# WTR: A Toolkit for Functional Anterograde Transneuronal Circuit Mapping

**DOI:** 10.64898/2026.03.23.713595

**Authors:** Chao Chen, Ruogu Liu, Aijia Yi-Luo, Xiaoyan Cao, Jitao Hu, Shikang Guan, Siyuan Chang, Xiaoli Cui, Wei Zhou, Fei Zhao, Chun-Teng Huang, Xin Duan, Lily Y. Jan, Tongfei A. Wang

## Abstract

The brain coordinates animal physiology and behavior via neuronal circuits. To understand and simulate brain functions, it is essential to delineate the synaptic connectivity between neurons. Transneuronal tracers serve as powerful tools for such purposes. In response to the demand for anterograde tracers for circuit mapping and functional interrogation, we developed WTR, a fusion protein of mammalian codon-optimized **<u>W</u>**GA, **<u>T</u>**EV-protease cleavage sequence, and **<u>R</u>**ecombinase. WTR expressed via AAV vectors in cell-type-specific starter neurons reaches their postsynaptic neurons and releases Cre/Flpo upon exposure to TEV-protease expressed in downstream neurons. Accompanied by Cre/Flpo-dependent expression of EGFP, GCaMP7s, or ChR2, the toolkit enables labeling, recording, or manipulation of downstream neurons. We utilized WTR to characterize downstream neurons of either glutamatergic or GABAergic neurons in the preoptic area of anterior hypothalamus for their differential actions in thermoregulation or stress responses, respectively. These results establish WTR as a versatile platform for functional anterograde circuit mapping.

**Significance Statement:** Anterograde transneuronal tools are essential for defining how genetically specified neurons influence downstream circuits, but existing approaches are limited by tracing efficiency, starter-cell specificity, or functional accessibility. We developed WTR, which shows higher transneuronal transfer efficiency than WGA-Cre, AAV1-Cre, and mWmC in our comparisons. WTR can be combined with Cre or Flpo driver lines to trace downstream circuits from genetically defined starter neurons. In addition, WTR enables expression of circuit-mapping tools such as GCaMPs, ChR2, neurotoxins, or even CRISPR tools in postsynaptic neurons, allowing anatomical tracing and functional interrogation of connected circuits within the same experimental framework.

## Introduction

Neurons are building blocks of the nervous system. They communicate through synaptic connections, processing information by integrating signals from various sources and producing responses that are sent to different downstream targets. Understanding the wiring diagrams of neural circuits and decoding the functional roles of inter-connected neurons will provide essential insights into brain function. Such an effort starts by tracing and visualizing the detailed morphology and connectivity of neurons. Labeling axons with genetically encoded fluorescent proteins allows the detection of neuronal synaptic projections throughout the brain (1). When combined with sparse labeling (2–4), tissue clearing (5), and whole-brain optical imaging (6), this approach enables high-resolution, three-dimensional reconstructions of single-neuron projection patterns, while providing little information about the downstream neurons.

As an orthogonal approach for intrinsic neuronal labeling *per se*, transneuronal tracers may reveal neuronal connectivity by labeling pre-synaptic (retrograde) or post-synaptic (anterograde) neurons. Identification of neurons that provide synaptic inputs to a brain region or to a specific neuronal type can be achieved by using retrograde transneuronal tracers, such as rabies virus (RV) (7–10) and pseudorabies virus (PRV) (11). One pioneer example is the application of the RV SAD-B19 strain for retrograde tracing (8). Genetically modified SAD-B19 is compatible with transgenic mouse lines that express Cre or Flpo recombinase in specific neuronal types, enabling cell-type-specific targeting and tracing (12). By integrating GCaMP or ChR2 into the Rabies payload, upstream neurons can be functionally characterized based on their neuronal activity or functional outputs (8, 12, 13). As a status quo, anterograde transneuronal tracing tools at the same level of resolution and versatility are still largely missing (14).

Anterograde transneuronal tracers aim to label postsynaptic neurons specifically and efficiently. For example, existing viral approaches for anterograde circuit mapping, including VSV-, HSV-, and YFV-based systems, enable tracing from starter neurons to downstream targets (15–18). However, these systems can present practical challenges, including neurotoxicity, large viral genomes, and the complexity of viral engineering. ATLAS (19) was recently established as a nanobody-based targeting agent against the extracellular N-terminus of the AMPA receptor subunit GluA1 for anterograde monosynaptic labeling. However, its application is restricted to circuits containing GluA1-expressing postsynaptic neurons, rendering this approach inaccessible to GluA1-negative circuits. The most widely used anterograde tracer is high-titer AAV1 carrying Cre or Flpo recombinase, which can transfer from infected neurons to their downstream targets to enable manipulation of connected circuits (20, 21). However, AAV1 shows retrograde uptake and lacks cell-type specificity in targeting starter neurons at the injection site.

We adopted a different strategy to generate a genetically encoded anterograde transneuronal tracing toolkit derived from wheat germ agglutinin (WGA), a lectin derived from *Triticum vulgaris* that is widely used in cell biology and neuroscience (22, 23). Native WGA exhibits both anterograde and retrograde tracing capability (24, 25). Gradinaru and colleagues have linked the wildtype WGA to Cre recombinase and expressed the fusion protein using AAV viral vectors (26). In combination with Cre-dependent expression of the optogenetic opsin ChR2 in downstream neurons, WGA-Cre enables not only labeling but also perturbation of downstream neurons. Our recent effort established an anterograde transneuronal tracer mWGA-mCherry (mWmC) that fused a mammalian codon-optimized WGA (mWGA) with mCherry (27). The fusion protein enabled fluorescent labeling in postsynaptic neurons. AAV-encoding made it compatible with widely available transgenic tools, such as existing Cre driver lines for different neuronal cell types, while enabling stable expression and long-term labeling over weeks or months. However, such a single-component AAV-encoded tracer is not adaptable to secondary payloads, such as GCaMP or ChR2 for the purpose of functional perturbations in postsynaptic cells.

Building on mWmC, we integrated an anterograde transneuronal functional circuit-mapping toolkit, termed WTR, which includes a mammalian codon-optimized WGA (mWGA2.0), a tobacco etch virus protease (TEVp) cleavage sequence (TEVcs) (28, 29), and Cre or Flpo recombinase as a single fusion protein. This re-engineered WTR enables efficient anterograde transneuronal transfer of the fusion protein, mainly restricted to first-order downstream neurons while minimizing retrograde spread. The second part of the toolkit involves TEVp, which is delivered into postsynaptic target regions. Consequently, the recombinase (Cre or Flpo) is released from the fusion protein upon TEV protease cleavage. This cleavage step is designed to increase recombination efficiency by separating Cre or Flpo from the WGA carrier, thereby potentially improving nuclear access and site-specific recombination. By uncoupling recombinase activity from continued carrier spread, TEVp can also help restrict recombinase-dependent labeling to the initially transduced postsynaptic neurons. In addition, expression of TEVp in Cre-negative local neurons (using a Cre-off design) provides a further safeguard against local circuit-mediated spread of WTR within the injection site, thereby improving starter-cell specificity. Thus, the TEVp module provides an additional layer of control over both recombination efficiency and labeling specificity. Meanwhile, a third AAV component encoding Cre- or Flpo-dependent EGFP, GCaMP7s, or ChR2 can also be introduced into the same area, enabling post-synaptic neuron-specific labeling, recording, or functional perturbation upon request (**Fig.6**).

Following validations of the WTR toolkit, we primarily focused on mapping the anterograde transneuronal targets of the mouse preoptic area of the anterior hypothalamus (POA). Previous studies implicated POA in regulating various physiological processes, such as thermoregulation, feeding, drinking, sleep, and innate behaviors (30). Yet a detailed wiring diagram linking POA neuronal subtypes to specific functions remains undefined. Previous studies indicated multiple candidate target regions, including potential direct innervation of neurons in the dorsomedial hypothalamus (DMH) (31–33) and periaqueductal gray (PAG) (34). We first delineated the differential circuits downstream of either glutamatergic POA (POA^Vglut2^) or GABAergic POA (POA^VGat^) neurons using the WTR toolkit. In combination with ChR2 or DREADDs, we demonstrated that DMH neurons innervated by POA^Vglut2^ are involved in thermoregulation. In contrast, PAG neurons innervated by POA^Vglut2^ promote anxiety-like behavior. In these studies, we showed that the WTR toolkit is highly adaptable to existing genetic-based approaches throughout the CNS. By combining circuit tracing and functional interrogation into one integrated toolkit, this experimental design provides an efficient platform for decoding neural circuits at both anatomical and functional levels.

## Results

### WTR exhibits predominantly anterograde spread with minimal retrograde labeling

Building on the mWmC design, we generated WTR, a codon-optimized fusion protein in which mWGA2.0 is linked to a TEV protease cleavage site (TEVcs) followed by either Cre or Flpo recombinase. We also generated Cre- or Flpo-dependent WTR vectors for genetically restricted induction in defined starter populations. The complete designs of the WTR constructs, including the Igκ leader sequence, 3×HA tag, WPRE, and bGH polyadenylation signal, are shown in **Fig. 1A**.

**Figure 1.**
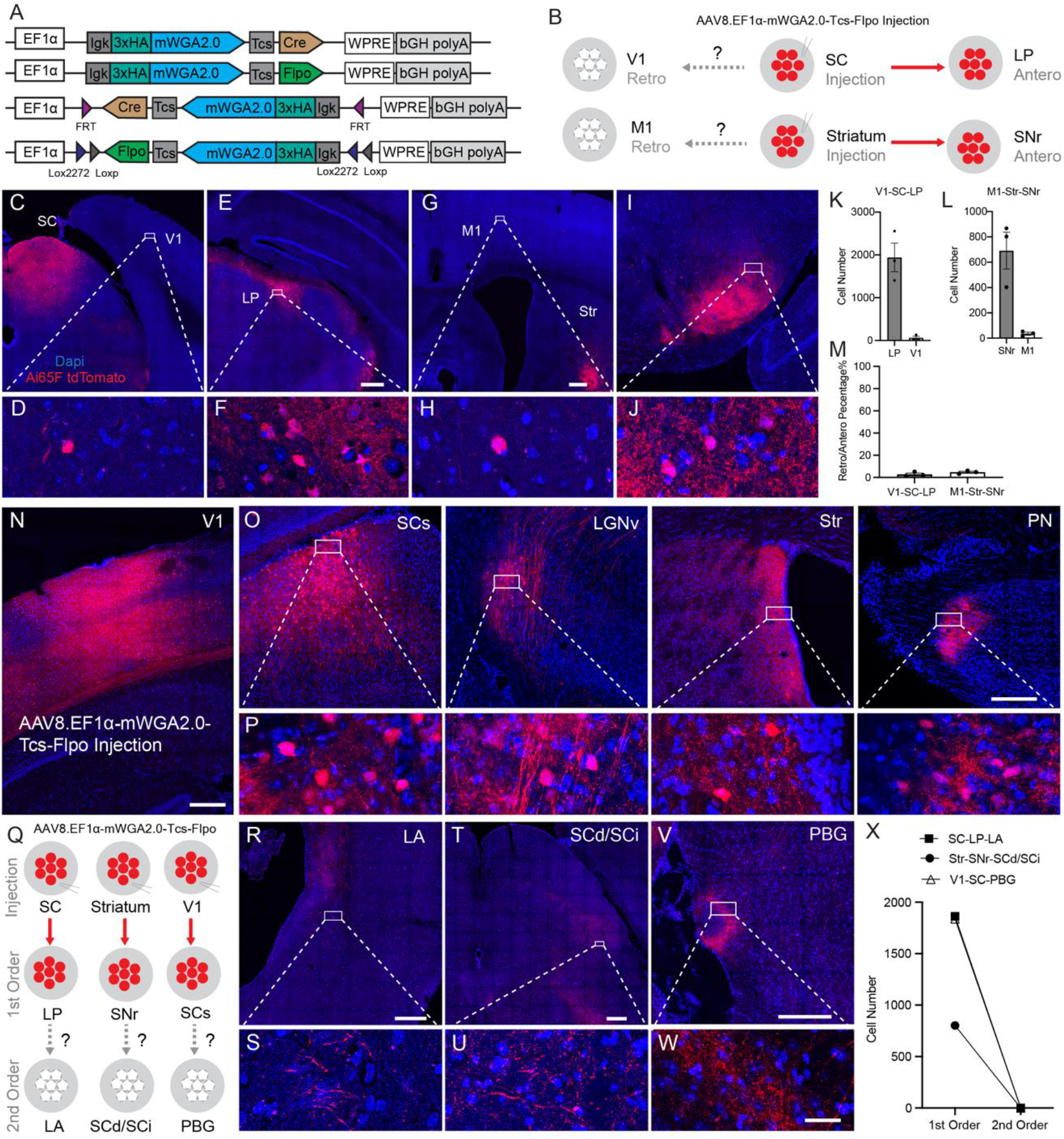
Directionality and first-order restriction of WTR transfer. **(A)** Schematic illustration of WTR constructs, including constitutive and Cre/Flpo-dependent designs. **(B)** Schematic illustration of the V1–SC–LP and M1–striatum–SNr circuits used to assess anterograde and retrograde transfer following WTR injection into SC or striatum. **(C-F)** Injection of AAV2/8.EF1α-mWGA2.0-TEVcs-Flpo into the superior colliculus (SC) of Ai65F mice and analysis of labeling in the V1–SC–LP pathway. Low-magnification image showing the SC injection site and V1 (**C**), with a corresponding high-magnification image of V1 (**D**), show few of retrograde V1 labeled cells, while robust anterograde labeling was detected in the lateral posterior thalamic nucleus (LP), a downstream target of SC (**E, F**). **(G-J)** Injection of AAV2/8.EF1α-mWGA2.0-TEVcs-Flpo into the striatum of Ai65F mice and analysis of labeling in the M1–striatum–SNr pathway. Low-magnification image showing the striatum injection site and M1 (**G**), with a corresponding high-magnification image of M1 (**H**) shows few of retrograde M1 labeled neurons, while anterograde labeling was highly detected in the substantia nigra pars reticulata (SNr) (**I, J**). **(K-M)** Quantification of labeled neurons in the anterograde and retrograde regions of the V1–SC–LP (**K**) and M1–striatum–SNr (**L**) pathways, with the corresponding retrograde to anterograde labeling percentages show in (**M**) (n = 3). **(N)** Representative image of the V1 injection site following injection of of AAV2/8.EF1α-mWGA2.0-TEVcs-Flpo into V1 of Ai65F mice. **(O, P),** Low (**O**) and high magnification (**P**) images showing WTR mediated labeling in first order downstream targets of V1, including the superficial superior colliculus (SCs), ventral lateral geniculate nucleus (LGNv), striatum (Str), and pontine nucleus (PN). tdTomato-positive neuronal somata were detected within the labeled axonal terminal fields in each first-order target region. **(Q)** Schematic illustration of three pathways used to examine potential second order WTR spread: SC–LP–LA, striatum–SNr–SCd/SCi, and V1–SCs–PBG. Solid red arrows indicate first-order projections, and dashed gray arrows indicate the corresponding potential second-order projections examined. **(R-W)** Analysis of potential second order labeling in the lateral amygdala (LA), deep/intermediate superior colliculus (SCd/SCi), and parabigeminal nucleus (PBG), downstream of LP, SNr, and SCs, respectively. Labeled axonal terminals were present in these regions, whereas tdTomato positive neuronal somata were not detected. Low magnification images are shown in **R**, **T**, and **V**, with corresponding high-magnification images in **S**, **U**, and **W**, respectively. **(X)** Quantification of number of labeled cells in the first and second order downstream regions in three different mice. No second-order labeled neuronal somata were detected at 2 months after injection. Red, tdTomato; blue, DAPI. Scale bars: 250 μm (**C, E, G, L, N, R, T, V**; the scale bar shown in **G** applies to **C**; the scale bar shown in **N** applies to **I**; the scale bar shown in the PN panel applies to all low-magnification panels in **O**); 25 μm (**D, F, H, J, P, S, U, W**; the scale bar shown in **W** applies to **D, F, H, J**, all high-magnification panels in **P**, and **S, U, W**).

A technical concern is the bidirectional transneuronal transfer properties of purified WGA proteins, especially when high concentrations of proteins exist at the axonal terminals (24, 25). AAV-encoded mWmC restricts protein production within a given cell and demonstrates minimal retrograde spread (27). WTR was built on the same framework; we first tested whether it likewise preserves strongly anterograde transfer with minimal retrograde labeling. The AAV2/8.EF1α-mWGA2.0-TEVcs-Flpo was injected into the superior colliculus (SC) of Rosa26-CAG-fDIO-tdTomato (Ai65F) mice (35), so that cells expressing or uptaking Flpo recombinase would express tdTomato (**Fig. 1C**). SC receives primary visual inputs from V1 cells and sends downstream projections to the lateral posterior thalamic nucleus (LP) (36, 37) (**Fig. 1B**). Four weeks after injection, robust labeling was observed in LP (**Fig. 1E,F**), whereas only a small number of tdTomato-positive neurons were detected in V1 (**Fig. 1C,D**). Quantification showed substantially more labeled neurons in LP than in V1 (**Fig. 1K**), yielding a low retrograde-to-anterograde labeling ratio (**Fig. 1M**). Notably, the downstream LP contained dense tdTomato-positive axonal fibers and terminals derived from the SC injection site, whereas such axonal labeling was absent in the upstream V1 region, consistent with the known direction of the SC-LP projection. Similarly, we examined the striatum, which receives excitatory inputs from the motor cortex (M1) and provides major inhibitory outputs to the substantia nigra pars reticulata (SNr) (38) (**Fig. 1B**). We injected AAV2/8.EF1α-mWGA2.0-TEVcs-Flpo into the striatum of Ai65F mice (**Fig. 1G**). We detected robust anterograde labeling in the SNr (**Fig. 1I, J**), whereas retrograde labeling in M1 was minimal (**Fig. 1G, H**). As in the SC–LP pathway, the downstream SNr contained prominent tdTomato-positive axonal labeling in addition to labeled neuronal somata, whereas the upstream M1 region showed little axonal signal, further supports the assignment of LP and SNr as downstream targets and V1 and M1 as upstream input regions in these experiments.

Given that dopaminergic neurons in the neighboring substantia nigra pars compacta (SNc) project to the striatum (39), we further examined whether the labeled somata observed in the substantia nigra could represent retrogradely labeled SNc neurons. Because tdTomato protein was present in both labeled neuronal somata and dense axonal projections, we combined tdTomato immunostaining with RNAscope detection of tdTomato and tyrosine hydroxylase (TH) mRNAs. tdTomato immunostaining provided the overall anatomical distribution of labeled fibers and cells, whereas tdTomato mRNA was used to identify locally labeled neuronal somata without interference from tdTomato-positive axons. The tdTomato mRNA-positive cells were TH-negative (**SI Appendix, Fig. S1**), supporting the conclusion that these labeled neurons were not dopaminergic SNc neurons. Together, these results show that WTR transfer is predominantly in the anterograde direction in multiple circuits.

### WTR traces predominantly first-order anterograde targets

We next tested whether WTR labeling extends beyond first-order downstream targets. We injected AAV2/8.EF1α-mWGA2.0-TEVcs-Flpo into the primary visual cortex (V1) of Ai65F mice (**Fig. 1N**). At 2 months after injection (mpi), tdTomato-expressing cell bodies were found interspersed within the terminal fields of tdTomato-labeled axons across all regions known to receive direct projections from V1 but have no projections to the V1 region (40–42), including SC, ventral lateral geniculate nucleus (LGNv), striatum, and pontine nucleus (PN) (**Fig. 1O,P**).

To determine whether WTR labeling continued beyond these first-order neurons, we examined three circuits in which the labeled first-order neurons project to neurons in known second-order regions (**Fig. 1Q**). In the lateral amygdala (LA) of the SC-LP-LA circuit, the deep/intermediate superior colliculus (SCd/SCi) of the striatum-SNr-SCd/Sci circuit, and the parabigeminal nucleus (PBG) of the V1-SCs-PBG circuit, we observed tdTomato-positive axonal terminals derived from labeled first-order neurons, but no tdTomato-positive neuronal somata (**Fig. 1R–W**). Thus, the first-order neurons were efficiently labeled to reveal their projections into these second-order regions, while there was no detectable transfer of WTR to the second-order neuronal somata. Quantification of labeled neuronal somata in the first- and second-order regions across all three pathways showed robust labeling in first-order targets but no detectable labeling of somata in the second-order regions examined (**Fig. 1X**). In summary, under our experimental conditions and up to 2 months after injection, WTR-mediated labeling was predominantly restricted to first-order downstream neurons, with no detectable second-order spread in the circuits examined.

### WTR exhibits higher anterograde transneuronal tracing efficiency

Having examined the directionality and first-order restriction of WTR transfer, we next compared its tracing efficiency with existing anterograde tools. We compared WTR against the original wildtype WGA-Cre (26), mWmC (27), and AAV2/1-Cre (20) (**Fig. 2A**). We also included a WTR condition without TEVp in the downstream DMH to directly assess the contribution of TEVp to overall tracing efficiency *in vivo*.

**Figure 2.**
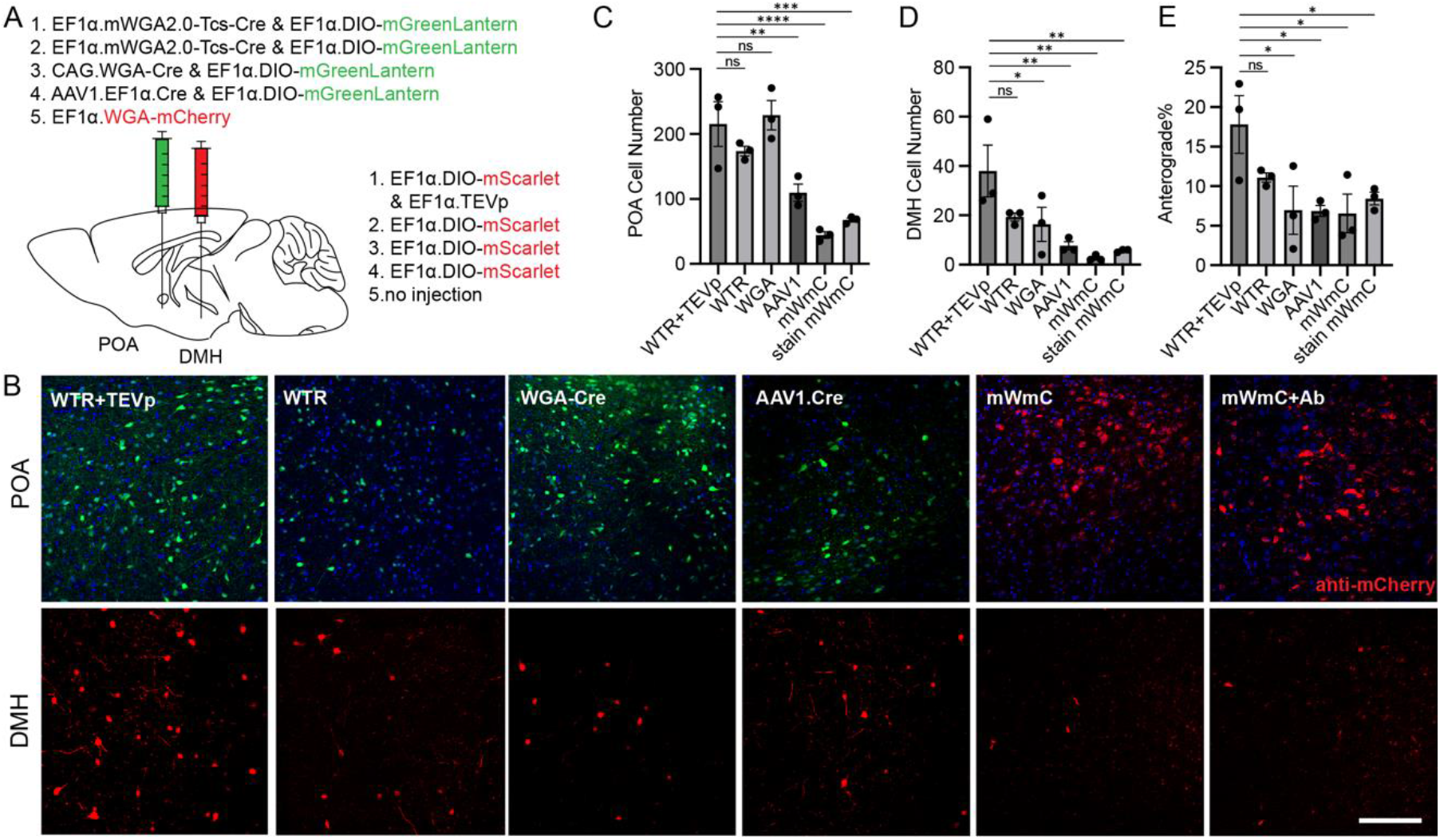
WTR exhibits higher anterograde tracing efficiency from POA to DMH, as compared to other viral tools **(A)** Viral injection setup for different anterograde tracing tools: **WTR+TEVp:** POA, AAV2/9.EF1α-mWGA2.0-TEVcs-Cre + AAV2/9.EF1α-DIO-mGreenLantern; DMH, AAV2/9.EF1α-DIO-mScarlet + AAV2/9.EF1α-TEVp; **WTR:** POA, AAV2/9.EF1α-mWGA2.0-TEVcs-Cre + AAV2/9.EF1α-DIO-mGreenLantern; DMH, AAV2/9.EF1α-DIO-mScarlet; **WGA-Cre:** POA, AAV2/9.CAG-WGA-Cre + AAV2/9.EF1α-DIO-mGreenLantern; DMH, AAV2/9.EF1α-DIO-mScarlet; **AAV1:** POA, AAV2/1.EF1α-Cre + AAV2/9.EF1α-DIO-mGreenLantern; DMH, AAV2/9.EF1α-DIO-mScarlet **mWmC:** POA, AAV2/9.EF1α-mWGA-mCherry; **mWmC + Ab:** POA, AAV2/9.EF1α-mWGA-mCherry, stained with anti-mCherry antibody. **(B)** Representative images showing fluorescent cells in POA and DMH. Scale bar: 100 μm.s **(C-E)** Quantification of labeled neurons in the POA (**C**) and downstream target DMH (**D**) for different groups, and corresponding anterograde tracing efficiency (**E,** DMH labeled cells / POA labeled cells, %). n=3; * p<0.05, ** p<0.01, *** p<0.001, one-way ANOVA followed by Tukey’s post-hoc test. Data are represented as mean ± SEM.

There are five conditions in these experiments. For the first two conditions, we injected AAV2/9.EF1α-mWGA2.0-TEVcs-Cre plus a Cre-dependent reporter (AAV2/9.EF1α-DIO-mGreenLantern) into POA, paired with injection of another Cre-dependent reporter (AAV2/9.EF1α-DIO-mScarlet) either with or without AAV2/9.EF1α-TEVp into DMH of the wild-type (WT) mice. DMH is a region known to receive direct projections from POA (31–33). The third and fourth ones were AAV2/9.CAG-WGA-Cre and AAV2/1.EF1α-Cre, plus the Cre-dependent reporter AAV2/9.EF1α-DIO-mGreenLantern, into POA, paired with injection of the Cre-dependent reporter AAV2/9.EF1α-DIO-mScarlet into the DMH of WT mice. Last, we also injected AAV2/9.EF1α-mWGA-mCherry into the POA of WT mice (**Fig. 2A**). At 3 wpi, Cre-dependent mScarlet (or mCherry for mWmC in the last set)-expressing neurons were detected in DMH (**Fig. 2B**), indicating anterograde transneuronal transfer of WTR or other anterograde tracers from POA neurons onto the postsynaptic targets. As stationary markers, the mGreenLantern-labeled neurons in the POA indicated the transfection efficiency of the viral vectors. The number of mScarlet (or mCherry for mWmC)-expressing neurons in DMH, normalized by the number of mGreenLantern (or mCherry for mWmC)-expressing neurons in POA, was used to evaluate anterograde tracing efficiency. Given that mCherry is less bright than mScarlet, we applied an anti-RFP antibody to amplify the mCherry signals. Compared with WGA-Cre, AAV2/1-Cre, and mWmC, WTR showed higher normalized anterograde tracing efficiency in the POA–DMH circuit (**Fig. 2C–E**). These results indicate that WTR provides more efficient anterograde transneuronal labeling than the existing tools tested under the same experimental framework.

Given the possibility that Cre and FLPo may differ in recombination efficiency (43), we next tested whether the performance of WTR depends on the recombinase used. We directly compared AAV2/9.EF1α-mWGA2.0-TEVcs-Cre and AAV2/9.EF1α-mWGA2.0-TEVcs-FLPo in the same POA–DMH circuit under matched experimental conditions. Both constructs produced similar labeling at the POA injection site and in downstream DMH neurons. Quantification showed no significant difference in the number of labeled starter neurons, labeled downstream neurons, or normalized anterograde tracing efficiency (**SI Appendix, Fig. S2**). These results indicate that WTR supports comparable anterograde tracing using either Cre or FLPo.

To compare the WTR anterograde efficiency among different AAV serotypes, we injected AAV vectors expressing WTR [AAV2/2.EF1α-mWGA2.0-TEVcs-Cre (Serotype 2), AAV2/8.EF1α-mWGA2.0-TEVcs-Cre (Serotype 8), or AAV2/9.EF1α-mWGA2.0-TEVcs-Cre (Serotype 9)] plus a Cre-dependent reporter (AAV2/9.EF1α-DIO-mScarlet) into POA.

In the downstream DMH region, we injected a mixture of AAV2/9.EF1α-DIO-mGreenLantern and AAV2/9.EF1α-TEVp (**SI Appendix, Fig. S3A**). All three serotypes tested (AAV2/2, AAV2/8, AAV2/9) showed comparable normalized anterograde tracing efficiency (**SI Appendix, Fig. S3B, C**). Notably, AAV2/9 exhibited the best transfection and expression efficiency in POA starter neurons, yielding the highest number of marked neurons in the downstream DMH (**SI Appendix, Fig. S3B, C**). We therefore used AAV2/9 for most of the subsequent experiments because of its higher transfection efficiency in hypothalamus, unless otherwise stated.

Because POA and DMH are reciprocally connected (31, 44), we further tested whether the WTR-labeled DMH neurons could instead represent neurons projecting back to POA. We combined WTR-mediated anterograde tracing with AAV2.retro-mediated retrograde tracing in the same animals (**SI Appendix, Fig. S4A**). AAV2.retro-mScarlet that was injected into POA labeled DMH neurons projecting to POA, whereas WTR-dependent mGreenLantern expression was detected in DMH neurons receiving projections from POA. Only minimal overlap was observed between the anterogradely labeled DMH neurons and the retrogradely labeled DMH neurons (**SI Appendix, Fig. S4B, C**). These results indicate that the majority of WTR labeled DMH neurons are distinct from DMH neurons that innervate POA neurons and further support the predominantly anterograde transfer of WTR in this reciprocally connected circuit.

### TEVp-mediated cleavage enhances functional recombinase activity

WTR differs from earlier tools not only by its mWGA2.0 backbone but also by the incorporation of a TEV protease cleavage module. We thus asked whether TEVp-mediated cleavage contributes to WTR performance by enhancing recombinase activity. We first asked whether TEVp enhances the functional activity of the recombinase carried by WTR. For this purpose, we performed an established dual-luciferase assay (45), that measures the Cre-driven luciferase expression in HEK cells expressing WTR with or without TEVp (**SI Appendix, Fig. S5A**). We found that the Cre recombinase cleaved from WTR produced significantly higher level of Gaussia luciferase luminescence, indicating enhanced catalytic efficiency, as compared with that of WTR without TEVp (**SI Appendix, Fig. S5B**). These data suggest that releasing Cre from the WGA carrier improves its functional recombination efficiency. We then compared WTR-mediated tracing *in vivo* with or without TEVp expression in the downstream DMH (**Fig. 2A–E**). Although the WTR+TEVp group showed a higher mean normalized tracing efficiency, the difference compared with WTR without TEVp was not statistically significant.

We next tested whether TEVp can efficiently cleave WTR *in vivo*. We injected AAV2/9.EF1α-TEVp into the upstream region (POA) either two weeks before or concurrently with an injection of AAV2/9.EF1α-mWGA2.0-TEVcs-Cre and AAV2/9.EF1α-DIO-mScarlet into the POA, which was performed at the same time as the injection of AAV2/9.EF1α-DIO-mGreenLantern and AAV2/9.EF1α-TEVp into DMH (**SI Appendix, Fig. S5C**). Brain tissue was harvested, sectioned, and imaged three weeks after the DMH injection (**SI Appendix, Fig. S5D**). Samples from mice with an injection of TEVp viral constructs two weeks prior to injection of WTR showed diminished anterograde tracing (as indicated by mGreenLantern expression in DMH) compared to samples from mice with the delivery of WTR without TEVp or with concurrent delivery of both to POA (**SI Appendix, Fig. S5E**). This result indicates that TEVp derived from viral delivery two weeks before the appearance of WTR provides sufficient TEVp catalytic activity to efficiently cleave the TEVcs site within WTR, thereby reducing Cre/Flpo transfer from those neurons with TEVp expression in the mouse brain.

### Cell-type-specific downstream labeling and functional readout with WTR

We have found WTR to be a high-efficiency anterograde tracer that delivers a recombinase payload, thereby making downstream neurons accessible for recording or manipulating their neuronal activity. In contrast to AAV1-Cre, which lacks intrinsic starter-cell specificity at the injection site, WTR can be combined with Cre- or Flpo-based strategies to initiate tracing from cell-type-specific neuronal populations. However, dense local connectivity within the starter region creates a potential source of nonspecific labeling. WTR released from genetically defined starter neurons may be transferred to nearby Cre-negative neurons through local synaptic connections, allowing these local recipient neurons to become secondary sources of WTR and thereby reducing starter-cell specificity. To limit this local circuit-mediated propagation, we introduced DO-TEVp, which is expressed in Cre-negative neurons in the same brain region that contains the starter neurons, to cleave WTR fusion protein and minimize the contamination from WTR transfer via local synaptic connection. In the Cre-positive starter neurons, DO-TEVp is not expressed, allowing the intact WTR to deliver recombinase to postsynaptic neurons (**Fig. 6B**). In downstream, TEVp cleavage releases Cre/Flpo from WTR to drive recombinase-dependent expression of reporters, sensors, or effectors, while limiting recombinase relay to other neurons.

To test whether DO-TEVp improves the specificity of downstream functional access, we injected AAV2/9.EF1α-DO-TEVp into the POA two weeks before delivering AAV2/9.EF1α-DIO-mWGA2.0-TEVcs-Flpo to the POA, together with AAV2/9.EF1α-fDIO-ChrimsonR-mScarlet for Flpo-dependent optogenetic activation via ChrimsonR. In the downstream DMH, we co-injected the Flpo-dependent Ca^2+^ sensor AAV2/9.EF1α-fDIO-GCaMP7s and AAV2/9.EF1α-TEVp in Vgat-Cre or Vglut2-Cre mice. Three weeks after injection, *ex vivo* Ca^2+^ imaging was performed using freshly prepared DMH brain slices (**Fig. 3A**). In response to red laser stimulation of ChrimsonR-expressing axon terminals (635 nm, 10 ms pulses, 20 Hz, 120s duration, 5 mW), intracellular Ca^2+^ increases or decreases were detected in DMH neurons receiving synaptic projections from POA (**Fig. 3B, C**). Neuronal responses were classified based on deviations exceeding three standard deviations (3σ) from baseline activity during laser stimulation. Cells showing increases above +3σ were defined as activated, while those showing decreases below −3σ were defined as inhibited. In Vglut2-Cre mice, laser-stimulation of POA axon terminals produced predominantly excitatory responses in WTR-labelled DMH neurons (88/361 activated, 16/361 inhibited, 257/361 unchanged, **Fig. 3B, E**). By contrast, in Vgat-Cre mice, responses were largely inhibitory (14/282 activated, 81/282 inhibited, 187/282 unchanged, **Fig. 3C, E**). These opposing response profiles are consistent with the expected sign of Vglut2 vs. Vgat neuronal inputs and support the notion that WTR enables cell-type-specific anterograde targeting of downstream neurons for functional readout.

**Figure 3.**
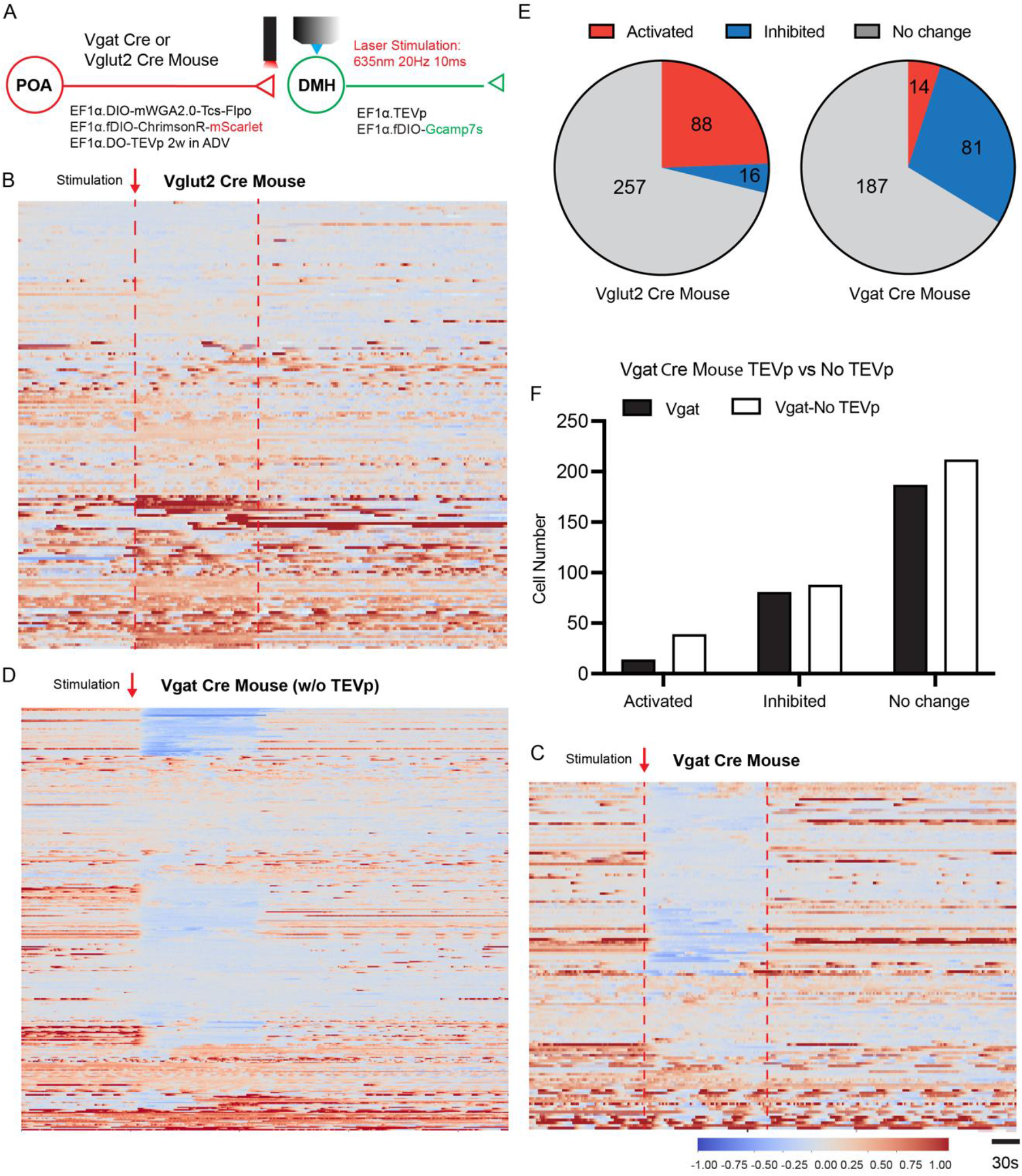
WTR enables recordings with downstream neurons of cell-type-specific starter neurons **(A)** Viral injection and experimental setup for Ca^2+^ imaging in DMH neurons. **(B-D)** Heatmaps of normalized z-scores showing the activity of DMH GCaMP7s-positive neurons (activated, inhibited, or no change) in response to red light stimulation (10 ms pulse, 20 Hz, 120 s duration, 5 mW) in Vglut2-Cre mice with TEVp injection (**B**), Vgat-Cre mice with TEVp injection (**C**), and Vgat-Cre mice without TEVp injection (**D**). **(E)** Pie chart showing the proportions of activated, inhibited, and unchanged responses across Vglut2-Cre mice and Vgat-Cre mice. **(F)** Cell counts of activated, inhibited, and unchanged DMH neurons in Vgat-Cre mice with or without TEVp Injection. The proportion of activated cells was compared between groups (39/339 cells without TEVp versus 14/282 cells with TEVp; n = 3 mice per group; p = 0.014, chi-square test).

To assess whether DO-TEVp injection into the upstream brain region contributes to this specificity, we repeated the Vgat-Cre experiment without DO-TEVp. We injected AAV2/9.EF1α-DIO-mWGA2.0-TEVcs-Flpo into the POA, paired with injection of Flpo-dependent AAV2/9.EF1α-fDIO-ChrimsonR-mScarlet plus AAV2/9.EF1α-fDIO-GCaMP7s into DMH of Vgat-Cre mice. These mice lacking TEVp in upstream Cre-negative cells showed 39/339 activated and 212/339 unchanged DMH neurons, compared with 14/282 activated and 187/282 unchanged DMH neurons in Vgat-Cre mice with DO-TEVp injection. In contrast, the number of inhibited neurons was comparable (88/339 versus 81/282) (**Fig. 3C, D, F**). Accordingly, the proportion of activated neurons was significantly higher without TEVp. These data indicate that application of Cre-off TEVp to the upstream brain region with starter neurons suppresses off-target recombinase transfer and/or local network-associated spread during anterograde tracing, thereby enhancing cell-type-specificity of the anterograde transneuronal tracing of WTR.

### Synaptic Specificity of Anterograde Spread of WTR

It is important to determine whether WTR spreads preferentially label synaptically connected target neurons, rather than being released from axons and then picked up by nearby cells. Electrophysiological recordings of post-synaptic currents, elicited by either electrical or optogenetic stimulation of presynaptic axon terminals, are considered the gold standard for confirming mono-synaptic transmission (46). We applied ChR2-assisted circuit mapping (CRACM) experiments to verify that WTR-labeled downstream neurons in DMH receive direct synaptic inputs from POA neurons. We first injected AAV2/9.EF1α-DO-TEVp into the POA of Vgat-Cre mice. Two weeks later, we injected Cre-driven AAV2/9.EF1α-DIO-mWGA2.0-TEVcs-Flpo plus AAV2/9.EF1α-fDIO-ChR2-EGFP into POA, paired with injection of the Flpo-dependent reporter AAV2/9.EF1α-fDIO-mScarlet and AAV2/9.EF1α-TEVp into DMH (**Fig. 4A**). Three weeks after the second injection, brain slices of DMH were prepared, and whole-cell patch-clamp recordings were performed on DMH neurons labeled with or without mScarlet. We performed voltage-clamp recordings of inhibitory post-synaptic currents (IPSCs), which were elicited by laser stimulation of ChR2-expressing axon terminals in DMH (460 nm, 5 ms pulses, 1 Hz). Blocking action potentials with tetrodotoxin (TTX, 1 μM) abolished the opto-evoked IPSCs, whereas enhancing neuronal excitability with 4-aminopyridine (4-AP, 1 mM) in the presence of TTX rescued only monosynaptic transmission. Representative sample traces classified as monosynaptic, polysynaptic, or no detectable synaptic transmission are shown in **Fig. 4B**. The amplitude of IPSC, especially after 4-AP treatment, provided an assessment of the monosynaptic tracing by WTR. The amplitudes of IPSCs across ACSF, ACSF+TTX, and ACSF+TTX+4-AP conditions were quantified separately in mScarlet^+^ and mScarlet^−^ DMH neurons (**Fig. 4C, D**). We determined that all the recorded mScarlet-positive neurons exhibited monosynaptic IPSCs (n = 14, median amplitude 275 pA, 100% reversible after 4-AP treatment), whereas only 4 of 15 mScarlet-negative neurons displayed monosynaptic IPSCs (median amplitude 97 pA, 100% reversible after 4-AP treatment); the remaining mScarlet-negative neurons exhibited either polysynaptic IPSCs (2 of 15, median amplitude 482 pA, irreversible after 4-AP treatment) or non-synaptic responses (9 of 14, amplitude too low to distinguish from noise) (**Fig. 4E, F**). This result suggests that WTR preferentially labels downstream DMH neurons that receive direct monosynaptic projections from POA Vgat neurons.

**Figure 4.**
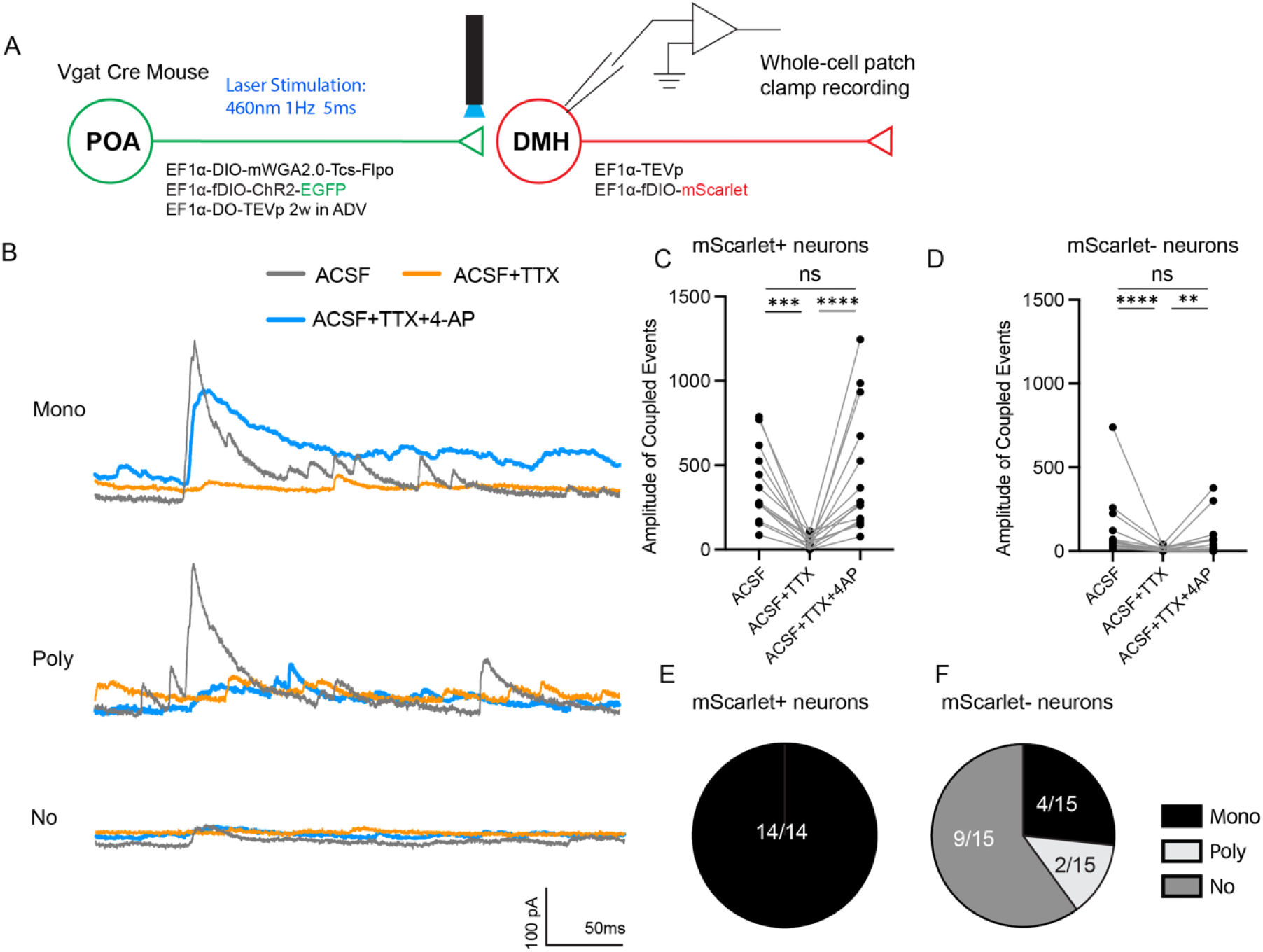
WTR labels downstream neurons that are monosynaptically connected with starter neurons **(A)** Viral injection strategy and experimental setup for whole-cell patch-clamp recordings from DMH neurons. **(B)** Representative traces showing monosynaptic, polysynaptic, or no evoked inhibitory postsynaptic currents (IPSCs) in DMH neurons during blue-light stimulation (460 nm, 5 ms pulses, 1 Hz, 5 mW) under ACSF, ACSF+TTX (1 μM), and ACSF+TTX (1 μM)+4-AP (1 mM) conditions. **(C,D)** IPSC amplitudes in mScarlet+ (**C**) and mScarlet− (**D**) DMH neurons under ACSF, ACSF+TTX, and ACSF+TTX+4-AP conditions. n = 14–15 cells for panels **C–F**. Amplitudes were compared using Friedman tests. **P < 0.01, ***P < 0.001, ****P < 0.0001. **(E,F)** Fraction of recorded mScarlet+ (**E**) and mScarlet− (**F**) DMH neurons exhibiting monosynaptic, polysynaptic, or no optogenetically evoked postsynaptic currents. Proportions were compared using chi-square tests; p = 0.0003.

### Manipulations of downstream neurons by WTR *in vivo*

Previous studies have shown that POA glutamatergic neurons drive hypothermic regulation of mouse physiology and behavior (47). To extend WTR from circuit mapping to functional interrogation *in vivo*, we asked whether WTR could enable projection-defined, cell-type-specific manipulation of downstream neurons in behavioral experiments. Given that thermoregulation is a physiological process with a relatively long time-course (a few hours), we first employed chemogenetic manipulation to investigate the POA-DMH pathway. We injected AAV2/9.EF1α-DO-TEVp into the POA two weeks before injection of AAV2/9.EF1α-DIO-mWGA2.0-TEVcs-Flpo into the POA, paired with injection of AAV2/9.EF1α-fDIO-hM3Dq-mGreenLantern (Flpo-dependent DREADD receptor) plus AAV2/9.EF1α-TEVp into DMH of Vglut2-Cre mice (**Fig. 5A**). Representative images confirmed hM3Dq-mGreenLantern expression in WTR-labeled DMH neurons downstream of POA^Vglut2^ or POA^Vgat^ neurons (**SI Appendix, Fig. S6A, B**). The body temperature of these mice was monitored using implantable temperature probes (**Fig. 5B**). Intraperitoneal (IP) administration of the DREADD agonist, deschloroclozapine (DCZ, 1 mg/kg), induced a significant decrease in the core body temperature (**Fig. 5C**). In contrast, using exactly the same strategy to activate the downstream DMH neurons of POA^Vgat^ neurons did not affect thermoregulation (**Fig. 5D**).

**Figure 5.**
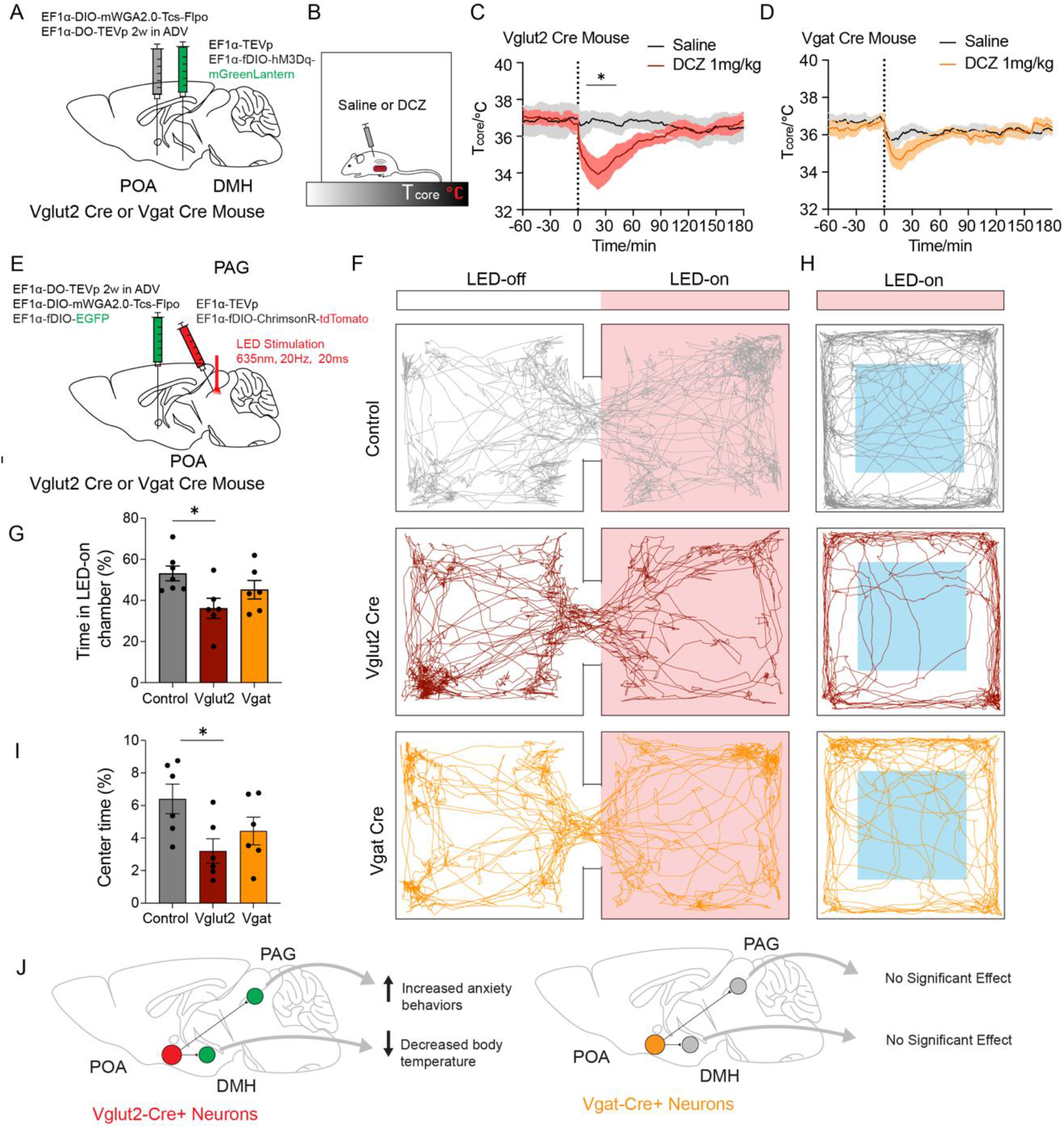
WTR enables functional manipulation of neurons downstream of cell-type-specific starter neurons **(A-B)** Schematic illustration of viral injections for chemogenetic manipulation of POA-innervated DMH neurons. T_core_ was measured using an implanted temperature probe. **(C-D)** DCZ (1 mg/kg; red and orange) or vehicle (saline; black) was injected intraperitoneally into Vglut2-Cre (**C**) or Vgat-Cre (**D**) mice expressing hM3Dq. Dashed lines indicate the time of injection. Vglut2-Cre: n=7, Interaction (F (240, 1446)=3.079, P<0.0001), drug (F (1, 1446)=340.4, p<0.0001), time (F (240, 1446)=0.7253, p=0.9991; Repeated measure two-way ANOVA with Šídák’s multiple comparisons test. Drug effect during 9-10, 37-39 min (P<0.05), 10–14, 33-36 min (P<0.01), 15-16, 27-32 min (P<0.001), and 17–26 min (P<0.0001) was significantly different.; Vgat-Cre: n=7, Interaction (F (240, 1446)=0.6256, p>0.9999), drug (F (1, 1446)=90.60, p<0.0001), time (F (240, 1446)=2.295, p<0.0001); Repeated measure two-way ANOVA with Šídák’s multiple comparisons test. Data are represented as mean ± SEM. **(E)** Schematic illustration of viral injection and stimulation setup (635nm, 10 Hz, 20 ms). **(F)** Representative locomotion traces for wildtype (gray), Vglut2-Cre (dark red), and Vgat-Cre (orange) animal groups in the two-chamber place preference test (PPT). **(G)** Quantification of the percentage of time spent in the LED-on chamber. n=6-7 animals per group, males only; *p<0.05, one-way ANOVA followed by Tukey’s post-hoc test. Data are represented as mean ± SEM. **(H)** Traces of locomotion for representative wildtype (gray), Vglut2-Cre (dark red), and Vgat-Cre (orange) animal groups during red light laser stimulation. Light blue squares indicate the designated center zone. **(I)** Quantification of the percentage of time spent in the center zone during the open field test (OFT). n=6 animals per group, males only; *p<0.05, one-way ANOVA followed by Tukey’s post-hoc test. Data are represented as mean ± SEM. **(J)** Summary schematic of projection-defined functional effects revealed by WTR. Manipulation of downstream neurons innervated by POA^Vglut2^ inputs decreased core body temperature via DMH targets and increased anxiety-like behaviors via PAG targets, whereas similar manipulations initiated from POA^Vgat^ inputs produced no significant effects.

**Figure 6.**
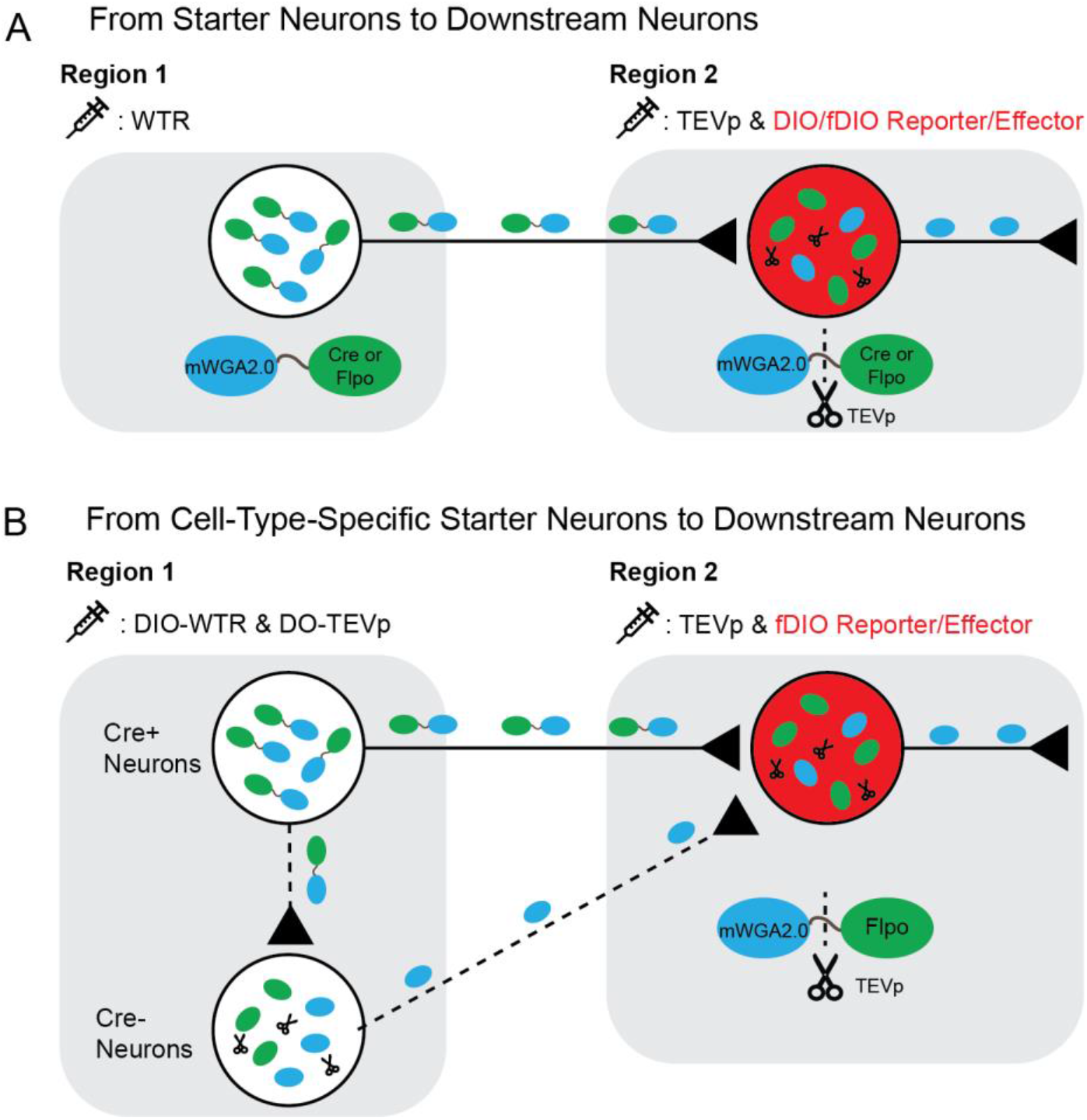
Schematic overview of WTR-based anterograde circuit tracing and functional access. **(A)** General WTR strategy. WTR, consisting of mWGA2.0 linked through a TEV protease cleavage site to Cre or Flpo recombinase, is expressed in starter neurons in Region 1 and transferred to downstream neurons in Region 2. TEVp expressed in the downstream region cleaves the recombinase from the WGA carrier, enhancing functional recombinase activity and enabling expression of Cre- or Flpo-dependent reporters, sensors, or effectors. **(B)** Cell-type-specific WTR strategy. In Cre-driver mice, Cre-dependent WTR (DIO-WTR) restricts WTR expression to Cre-positive starter neurons. A Cre-off TEVp construct (DO-TEVp) expresses TEVp in neighboring Cre-negative neurons, providing an additional safeguard against local circuit-mediated spread of WTR within the starter region. WTR produced by Cre-positive starter neurons is transferred to downstream neurons, where TEVp cleavage releases Flpo and activates Flpo-dependent reporters, sensors, or effectors. This configuration enables functional access to downstream neurons originating from genetically defined starter populations.

While chemogenetics is ideal for studying slow physiological processes such as thermoregulation, we further expanded WTR to include optogenetic modalities *in vivo*. We targeted the POA to PAG pathway, which is reported to modulate rapid behavioral shifts such as negative valence and anxiety-like behaviors (34). We injected AAV2/9.EF1α-DO-TEVp into the POA two weeks before injection of AAV2/9.EF1α-DIO-mWGA2.0-TEVcs-Flpo and AAV2/9.EF1α-fDIO-EGFP into the POA, paired with injection of AAV2/5.EF1α-fDIO-ChrimsonR-tdTomato plus AAV2/9.EF1α-TEVp into the PAG of WT mice, Vglut2-Cre mice, or Vgat-Cre mice (**Fig. 5E**). Representative images confirmed ChrimsonR-tdTomato expression in WTR-labeled PAG neurons downstream of POA^Vglut2^ or POA^Vgat^ neurons (**SI Appendix, Fig. S6C, D**). In the place preference test, LED stimulation was applied whenever the animal entered the designated stimulation (LED-ON) chamber (**Fig. 5F**). The animals with ChrimsonR-expressing PAG neurons that receive POA^Vglut2^ innervation spent less time in the LED-ON chamber as compared with WT animals (**Fig. 5G**). In contrast, similar experiments with the Vgat-Cre mice show no significant difference (**Fig. 5G**). In the open field test, optogenetic stimulation of ChrimsonR-expressing PAG neurons that receive projections from POA^Vglut2^ neurons spent significantly less time in the center zone compared to WT animals that were also exposed to light (**Fig. 5H, I**). In contrast, similar experiments with Vgat-Cre mice showed no significant difference in time spent in the center zone (**Fig. 5I**). A schematic summary of these projection-defined functional effects is shown in **Fig. 5J**. Together, these results demonstrate that WTR can be used to functionally manipulate neurons in downstream target regions that receive projections from specific types of neurons in the upstream brain regions.

## Discussion

In this study, we re-engineered WTR, an anterograde transneuronal system for functional circuit mapping. Across multiple CNS circuits, WTR demonstrated efficient, strongly biased anterograde spread, minimal retrograde labeling, and limited second-order transmission. WTR tracing can be initiated from genetically defined starter neurons and combined with recombinase-dependent functional payloads, such as Ca^2+^ indicators, chemogenetics, or optogenetic effectors. This complementary toolbox enabled circuit mapping and rapid functional validation within a single cohort of animals, thereby efficiently and comprehensively establishing links among synaptic connectivity, neuronal physiology, and animal behavior.

The original WGA protein exhibited both anterograde and retrograde spread (22, 23). This concern could mitigate the findings, especially as retrograde transfer may occur via axonal uptake of WGA protein at the injection site rather than via retrograde transport of WGA from starter cells (25). Similar to the mWmC design, WTR showed predominantly anterograde transfer in both the V1–SC–LP and M1–striatum–SNr pathways, while a small number of retrogradely labeled neurons were also detected. We therefore describe WTR as exhibiting predominantly anterograde transfer with minimal retrograde labeling. (**Fig. 1**). An important difference between WTR and direct application of WGA protein is that WTR is genetically expressed only in the starter neurons. This limits direct uptake of extracellular WGA by afferent axon terminals at the injection site, although it does not completely exclude retrograde transfer after WTR release. For applications in which residual retrograde labeling remains a concern, WTR can be combined with an AAV2.retro tracer injected into the starter region to identify neurons that project back to the starter population and distinguish them from predominantly anterogradely labeled downstream neurons, as demonstrated in the POA–DMH circuit (**SI Appendis, Fig. S4**). The cellular mechanism underlying the strong anterograde bias of genetically expressed WTR remains an open question.

Another important feature of anterograde tracers is the extent of higher-order spread. While we have demonstrated the potential application of WTR for labeling downstream neurons, its design does not preclude further relay beyond first-order targets. To directly assess this possibility, we examined several potential second-order targets and detected labeled axonal terminals but no labeled neuronal somata in the tested second-order regions (**Fig. 1**). These results indicate that WTR labeling is largely confined to first-order targets under the conditions examined.

A third key question is whether WTR transfer reflects true synaptic connectivity rather than local, nonspecific tracer uptake. Our electrophysiological validation provides direct evidence that WTR-labelled postsynaptic neurons are synaptically connected with downstream targets (**Fig. 4**). In all WTR-labelled (mScarlet^+^) DMH neurons tested (14/14; 100%), monosynaptic IPSCs were observed. Therefore, WTR labeling marks monosynaptically connected postsynaptic neurons, supporting synapse-dependent anterograde transfer. We recognize that the tracing efficiency of WTR does not reach 100%, as we detected a small percentage of DMH neurons that are not labeled by WTR from the POA but are nonetheless monosynaptic downstream neurons of POA neurons (**Fig. 4F**). Available transneuronal tracing tools do not label every anatomically connected neuron, and WTR likewise does not achieve 100% tracing efficiency.

Regarding the transneuronal efficiency, though it is hard to define the “ground truth”, we demonstrated that the WTR system exhibited higher transneuronal transfer efficiency as compared to existing tools such as WGA-Cre in a defined circuit (POA→DMH) (**Fig. 2**). One factor contributing to the improved tracing efficiency may be TEVp-mediated release of Cre/Flpo from WTR, which increased functional recombinase activity in our luciferase assay (**SI Appendix, Fig. S5**). However, direct comparison of WTR with and without TEVp in the downstream DMH showed no significant difference in the overall number of labeled downstream neurons or normalized tracing efficiency. This apparent difference between the biochemical assay and the *in vivo* tracing experiment may reflect the fact that the total reporter-positive cell number is influenced by both recombinase activity and the extent of WTR transfer. TEVp-mediated cleavage may increase the recombination efficiency of WTR delivered to a postsynaptic neuron while simultaneously limiting continued transfer of the intact WTR fusion protein to additional neurons. In the absence of TEVp, reduced recombinase activity could therefore be partly offset by increased local or secondary spread of intact WTR, resulting in a similar overall number of reporter-positive cells.

In contrast to AAV1-Cre, which lacks intrinsic starter-cell specificity at the injection site and therefore cannot initiate tracing from genetically defined populations, the Cre-dependent mWGA2.0-TEVcs-Flpo AAV vector enables tracing from cell-type-specific starter neurons in mouse lines with a Cre driver (**Fig. 3**). This application was further enhanced with TEVp expression in Cre-negative neurons (DO-TEVp, Cre-Off version) two weeks prior to the WTR expression at the injection site (**Fig. 6**). This procedure is designed to minimize the impact of local spread of WTR from Cre-positive neurons to Cre-negative neurons. Using cell-type-specific Ca^2+^ imaging *ex vivo*, we demonstrated that the WTR-labeled downstream neurons exhibit physiological responses largely aligned with the neurotransmitter identity of the upstream starter cells. Specifically, optogenetic activation of POA^Vglut2^ nerve terminals elicited predominantly excitatory Ca^2+^ transients in labeled DMH neurons, whereas optogenetic activation of POA^Vgat^ nerve terminals evoked largely inhibitory responses (**Fig. 3E**). However, a substantial fraction of WTR labeled neurons did not show detectable Ca²⁺ responses. Some synaptic inputs may be too weak to produce a detectable GCaMP7s signal, and some connections, particularly those located on distal dendrites, may be lost during brain-slice preparation. We also observed a small fraction of responses opposite to the expected transmitter identity. These responses may partly result from off-target expression at the injection site, as such unexpected responses increased when DO-TEVp was omitted. In addition, local polysynaptic circuits within the slice, including disinhibition or di-synaptic excitation, may also contribute to these responses. Therefore, the Ca²⁺ imaging experiments provide a population-level functional readout of WTR-labeled neurons but should not be interpreted as a direct test of monosynaptic connectivity for every labeled cell.

The WTR-based system enables anterograde transneuronal functional interrogation of downstream neurons that are difficult to accomplish with other systems like HSV, mainly due to viral toxicity that often limits experiments. This marks a significant technical advancement over many existing tools, such as mWmC, which only enables fluorescent labeling without functional testing. Using this WTR design paired with chemogenetics and optogenetic tools, we identified the downstream neurons of POA glutamatergic or GABAergic neurons in DMH or PAG. We used patch-clamp recordings to validate monosynaptic connectivity from POA starter neurons to WTR-labeled downstream neurons, while Ca²⁺ imaging was used to examine their functional responses to activation of upstream axons (**Figs. 3, 4**). We also manipulated the activity of these neurons and linked it to animal physiology and behavior (**Fig. 5**). However, the different effects observed after manipulating POA^Vglut2^ versus POA^Vgat^ downstream neurons should be interpreted with caution. Although we confirmed expression of hM3Dq or ChrimsonR in the corresponding DMH and PAG populations (**SI Appendix, Fig. S6**), the present experiments cannot determine whether the differences in physiological and behavioral effects result from differences in the number of labeled neurons or from distinct downstream neuronal populations with different functions. In future work, transcriptomic approaches such as single-cell RNA-sequencing could be used to define the molecular identities of WTR-labeled DMH and PAG neurons and determine whether different POA starter populations target distinct downstream cell types.

Finally, modular design of AAV-based WTR promotes rapid iteration for tool development: the vectors can be easily re-engineered and readily packaged into different AAV serotypes to suit diverse experimental scenarios. This engineering flexibility is superior to other viral tracers (HSV, VSV and YFV), where changes often require substantial genome re-engineering and re-optimization of toxicity. Such modularity should facilitate adaptation of WTR to a variety of circuits and potentially to peripheral nervous system applications.

In summary, the WTR is an effective tool when combined with prior anatomical knowledge to design WTR injections into the brain region containing upstream neurons and TEVp viral construct injections into the brain region containing putative downstream neurons, for anterograde tracing from specific neuronal types. Future applications of this toolkit include characterizing the transcriptome of these neurons using single-cell RNA-sequencing or spatial transcriptomics to identify their genetic markers, or perturbing their gene expression using CRISPR. This strategy can be applied to dissect neural circuits across a variety of brain regions involved in diverse physiological and pathological processes, making the potential applications of WTR and its derived tools broad and promising.

## Materials and Methods

| REAGENT or RESOURCE | SOURCE | IDENTIFIER |
| --- | --- | --- |
| **Antibodies** |  |  |
| goat anti-WGA | Vector Labs | AS-2024-1 |
| rabbit anti-mCherry | Abcam | ab167453 |
| **Bacterial and Virus Strains** |  |  |
| Adeno-associated virus-various | CIBR Vector Core and<br>Duan Lab @UCSF | N/A |
| **Chemicals, Peptides, and Recombinant Proteins** |  |  |
| Tetrodotoxin citrate | Tocris | 1069 |
| 4-Aminopyridine | MCE | HY-B0604 |
| Deschloroclozapine | MCE | HY-42110A |
| **Critical Commercial Assays** |  |  |
| ClonExpress Ultra One Step Cloning<br>Kit | Vazyme | C115 |
| Cypridina Luciferase Glow Assay Kit | Thermo Scientific | 16170 |
| Gaussia Luciferase Glow Assay Kit | Thermo Scientific | 16160 |
| RNAscope® Multiplex Fluorescent<br>Reagent Kit v2 | ACDBio | 323100 |
| TH-C2 | ACDBio | 317621-C2 |
| tdTomato-C3 | ACDBio | 317041-C3 |
| **Recombinant DNA** |  |  |
| pAAV-EF1α-mWGA2.0-TEVcs-Cre | This paper | |
| pAAV-EF1α-mWGA2.0-TEVcs-Flpo | This paper | |
| pAAV-EF1α-DIO-mWGA2.0-TEVcs-Flpo | This paper | |
| pAAV-EF1α-TEVp | This paper | |
| pAAV-DO-EF1α-TEVp | This paper | |
| pAAV-EF1α-TEVp-P2A-EGFP | This paper | |
| pAAV-EF1α-EGFP | This paper | |
| pBR322-EF1α-DIO-hGLuc-CMV-CLuc | This paper | |
| pAAV-CAG-WGA-Cre | Duan Lab @ UCSF |  |
| pAAV-EF1α-mWGA-mCherry | Duan Lab @ UCSF | |
| pAAV-EF1α-DIO-mScarlet | CIBR Vector Core | |
| pAAV-EF1α-DIO-mGreenLantern | CIBR Vector Core | |
| pAAV-EF1α-mScarlet | CIBR Vector Core | |
| pAAV-EF1α-DIO-GcaMP7s | CIBR Vector Core | |
| pAAV-EF1α-DIO-ChrimsonR-mScarlet | CIBR Vector Core | |
| pAAV-EF1α-fDIO-hM3Dq-mGreenLantern | CIBR Vector Core | |
| pAAV-EF1α-DIO-ChR2-EGFP | CIBR Vector Core | |
| pAAV-EF1α-DIO-mTagBFP2 | CIBR Vector Core | |
| pAAV-EF1α-fDIO-Chrimson-tdTomato | Addgene | 171027 |
| pAAV-EF1α-DIO-EYFP | Addgene | 27056 |
| **Experimental Models: Organisms/Strains** |  |  |
| Mouse: C57BL/6J | The Jackson<br>Laboratory | Stock#: 000664;<br>RRID:<br>IMSR_JAX:000664 |
| Mouse: Vglut2-Cre | The Jackson<br>Laboratory | Stock#: 016963;<br>RRID:<br>IMSR_JAX:016963 |
| Mouse: Vgat-Cre | The Jackson<br>Laboratory | Stock#: 028862;<br>RRID:<br>IMSR_JAX:028862 |
| Mouse: Rosa-LSL-H2B-GFP | The Jackson<br>Laboratory | Stock#: 036761;<br>RRID:<br>IMSR_JAX:036761 |
| **Software and Algorithms** |  |  |
| Prism 10.1.1 | Graphpad |  |
| VitalView | Starr Life Sciences |  |
| MouBeAT | Github |  |
| Fiji (ImageJ) | NIH |  |
| Illustrator 2024 | Adobe |  |
| Cellpose 2.0 | Carsen Stringer,<br>Marius Pachitariu | https://www.cellpose.org/ |
| RWD Fiber Photometry System | RWD Life Science |  |

### Animals

All procedures involving mice were approved by the Institutional Animal Care and Use Committee (IACUC) at the Chinese Institute for Brain Research in Beijing and at the University of California, San Francisco. Mice were maintained on a 12/12-hour light/dark cycle with regular chow and water provided *ad libitum*. The Vglut2-Cre, Vgat-Cre, and Ai65F transgenic mice were originally obtained from The Jackson Laboratory (stock numbers: 016963, 028862, and 032864, respectively) and were further bred at CIBR or UCSF animal facilities.

### Viral vectors

All plasmids were constructed using the ClonExpress Ultra One-Step Cloning Kit (Vazyme, Nanjing, China). The TEVp fragment was obtained from plasmid pAAV-flex-taCasp3-TEVp (Addgene, #45580) and used to replace Flpo in pAAV-EF1α-Flpo (Addgene, #55637), resulting in pAAV-EF1α-TEVp. In the Cre-dependent cassette (Flex-switch), TEVp was inserted in the 5’ to 3’ orientation to create pAAV-EF1α-DO-TEVp, enabling CreOFF-dependent control. EF1α-DIO-hGluc-CMV-CLuc was made from pC0037 (Addgene, #181934).

The codon-optimized WGA (mWGA) sequence was synthesized by Genewiz Inc. (South Plainfield, NJ) and fused with a TEVp cleavage site (GAGAACCTGTACTTCCAGGGC), along with either Cre or Flpo recombinase. The mWGA2.0-TEVcs-Flpo sequence was inserted in the 3’ to 5’ orientation to generate pAAV-EF1α-DIO-mWGA2.0-TEVcs-Flpo. The mWGA2.0-TEVcs-Cre and mWGA2.0-TEVcs-Flpo fragments were cloned into the pAAV-EF1α-DIO-mWGA-mCherry-T2A-YFP backbone (gift from the Xin Duan Lab) by replacing the original ORF. The complete designs of the WTR constructs are shown in Fig. 1a. The reference sequence for mWGA2.0-TEVcs-Flpo and mWGA2.0-TEVcs-Cre were as follows:

mWGA2.0-TEVcs-Flpo:

ATGGAGACAGACACACTCCTGCTATGGGTACTGCTGCTCTGGGTTCCAGGTTCCACTGGTGACTATCCATATGATGTTCCAGATTATGCGTCCGGCTATCCCTATGACGTCCCGGACTATGCAGGATCCTATCCATATGACGTTCCAGATTACGCAGGACCGGTCATGACAGCACAAGCCCAGAGGTGCGGCGAACAAGGCAGCAATATGGAATGTCCAAACAACCTCTGCTGCTCTCAGTATGGCTACTGCGGTATGGGAGGAGATTACTGTGGCAAGGGGTGCCAGAATGGTGCATGCTGGACATCCAAGCGATGCGGATCTCAGGCTGGGGGAGCAACCTGTCCTAACAACCATTGTTGTTCACAGTACGGACACTGTGGGTTTGGCGCAGAATACTGTGGTGCCGGATGCCAGGGCGGACCATGTAGAGCTGACATTAAGTGTGGCTCACAGTCTGGGGGTAAACTCTGCCCCAACAATCTTTGTTGTAGTCAGTGGGGCTTCTGTGGCCTCGGCTCCGAGTTTTGCGGCGGTGGATGCCAGTCCGGTGCATGCAGTACAGACAAGCCATGCGGGAAAGACGCAGGGGGCAGGGTGTGTACCAACAACTACTGTTGCTCCAAGTGGGGGTCATGTGGGATCGGCCCTGGCTATTGTGGGGCCGGATGTCAGTCAGGTGGATGTGATGCCGCCCGGGATCCACCGGTAGCATCCGCCACCGAGAACCTGTACTTCCAGGGCATGAGCCAGTTCGACATCCTGTGCAAGACCCCCCCCAAGGTGCTGGTGCGGCAGTTCGTGGAGAGATTCGAGAGGCCCAGCGGCGAGAAGATCGCCAGCTGTGCCGCCGAGCTGACCTACCTGTGCTGGATGATCACCCACAACGGCACCGCCATCAAGAGGGCCACCTTCATGAGCTACAACACCATCATCAGCAACAGCCTGAGCTTCGACATCGTGAACAAGAGCCTGCAGTTCAAGTACAAGACCCAGAAGGCCACCATCCTGGAGGCCAGCCTGAAGAAGCTGATCCCCGCCTGGGAGTTCACCATCATCCCTTACAACGGCCAGAAGCACCAGAGCGACATCACCGACATCGTGTCCAGCCTGCAGCTGCAGTTCGAGAGCAGCGAGGAGGCCGACAAGGGCAACAGCCACAGCAAGAAGATGCTGAAGGCCCTGCTGTCCGAGGGCGAGAGCATCTGGGAGATCACCGAGAAGATCCTGAACAGCTTCGAGTACACCAGCAGGTTCACCAAGACCAAGACCCTGTACCAGTTCCTGTTCCTGGCCACATTCATCAACTGCGGCAGGTTCAGCGACA TCAAGAACGTGGACCCCAAGAGCTTCAAGCTGGTGCAGAACAAGTACCTGGGCGTGATCATTCAGTGCCTGGTGACCGAGACCAAGACAAGCGTGTCCAGGCACATCTACTTTTTCAGCGCCAGAGGCAGGATCGACCCCCTGGTGTACCTGGACGAGTTCCTGAGGAACAGCGAGCCCGTGCTGAAGAGAGTGAACAGGACCGGCAACAGCAGCAGCAACAAGCAGGAGTACCAGCTGCTGAAGGACAACCTGGTGCGCAGCTACAACAAGGCCCTGAAGAAGAACGCCCCCTACCCCATCTTCGCTATCAAGAACGGCCCTAAGAGCCACATCGGCAGGCACCTGATGACCAGCTTTCTGAGCATGAAGGGCCTGACCGAGCTGACAAACGTGGTGGGCAACTGGAGCGACAAGAGGGCCTCCGCCGTGGCCAGGACCACCTACACCCACCAGATCACCGCCATCCCCGACCACTACTTCGCCCTGGTGTCCAGGTACTACGCCTACGACCCCATCAGCAAGGAGATGATCGCCCTGAAGGACGAGACCAACCCCATCGAGGAGTGGCAGCACATCGAGCAGCTGAAGGGCAGCGCCGAGGGCAGCATCAGATACCCCGCCTGGAACGGCATCATCAGCCAGGAGGTGCTGGACTACCTGAGCAGCTACATCAACAGGCGGATCTGA

mWGA2.0-TEVcs-Cre:

ATGGAGACAGACACACTCCTGCTATGGGTACTGCTGCTCTGGGTTCCAGGTTCCACTGGTGACTATCCATATGATGTTCCAGATTATGCGTCCGGCTATCCCTATGACGTCCCGGACTATGCAGGATCCTATCCATATGACGTTCCAGATTACGCAGGACCGGTCATGACAGCACAAGCCCAGAGGTGCGGCGAACAAGGCAGCAATATGGAATGTCCAAACAACCTCTGCTGCTCTCAGTATGGCTACTGCGGTATGGGAGGAGATTACTGTGGCAAGGGGTGCCAGAATGGTGCATGCTGGACATCCAAGCGATGCGGATCTCAGGCTGGGGGAGCAACCTGTCCTAACAACCATTGTTGTTCACAGTACGGACACTGTGGGTTTGGCGCAGAATACTGTGGTGCCGGATGCCAGGGCGGACCATGTAGAGCTGACATTAAGTGTGGCTCACAGTCTGGGGGTAAACTCTGCCCCAACAATCTTTGTTGTAGTCAGTGGGGCTTCTGTGGCCTCGGCTCCGAGTTTTGCGGCGGTGGATGCCAGTCCGGTGCATGCAGTACAGACAAGCCATGCGGGAAAGACGCAGGGGGCAGGGTGTGTACCAACA

ACTACTGTTGCTCCAAGTGGGGGTCATGTGGGATCGGCCCTGGCTATTGTGGGGCCGGATGTCAGTCAGGTGGATGTGATGCCGCCCGGGATCCACCGGTAGCATCCGCCACCGAGAACCTGTACTTCCAGGGCTCCAATCTCCTGACTGTTCACCAGAACCTCCCTGCGCTGCCAGTAGATGCCACTAGCGATGAGGTCAGGAAAAATCTCATGGATATGTTTAGGGATAGACAGGCGTTTTCTGAACACACCTGGAAAATGCTGCTTAGCGTGTGCCGATCCTGGGCAGCCTGGTGTAAGCTGAACAATCGCAAATGGTTCCCCGCCGAGCCGGAGGACGTGCGCGATTACCTGCTGTATCTCCAGGCAAGAGGGCTGGCTGTCAAGACTATCCAGCAGCACTTGGGCCAACTGAATATGCTGCATCGACGCAGCGGGCTCCCCCGGCCTAGCGATTCAAACGCAGTCTCCCTTGTTATGAGGAGAATTAGAAAGGAAAACGTAGATGCGGGTGAGAGGGCTAAGCAGGCTCTCGCTTTTGAGCGGACTGATTTCGACCAGGTCAGATCCCTGATGGAGAACAGCGATCGGTGCCAGGACATCAGGAACCTCGCATTTCTGGGAATTGCATATAACACACTTCTGCGCATAGCTGAGATCGCCCGGATCAGAGTGAAAGACATCAGTCGAACGGACGGCGGCCGGATGCTTATTCATATTGGACGCACAAAGACATTGGTCAGCACCGCTGGCGTTGAAAAGGCCTTGTCCCTGGGCGTAACGAAGCTGGTGGAAAGATGGATCTCAGTGTCCGGCGTGGCTGACGACCCTAATAATTACTTGTTCTGTCGAGTGAGAAAAAACGGAGTCGCCGCGCCCTCTGCCACCAGCCAATTGAGTACACGGGCCCTTGAAGGGATCTTTGAGGCAACCCACCGACTCATATACGGAGCCAAGGATGACAGTGGCCAGAGGTATCTCGCCTGGTCAGGTCATTCTGCTAGGGTGGGGGCCGCACGAGACATGGCGCGGGCAGGAGTCTCCATACCAGAGATTATGCAAGCTGGAGGTTGGACAAATGTGAACATCGTTATGAACTATATCCGCAATCTTGACTCTGAAACCGGGGCCATGGTGAGACTGCTCGAAGATGGTGACTG A

### Luciferase assay

In a 48-well plate, 250 ng of either the pAAV-EF1α-TEVp-P2A-EGFP expression vector or a control pAAV-EF1α-EGFP vector was transfected into HEK293T cells. After 24 hours, 125 ng of the pAAV-EF1α-mWGA2.0-TevCS-Cre expression plasmid and 125 ng of the dual luciferase reporter pBR322-EF1α-DIO-hGLuc-CMV-CLuc were co-transfected into the cells. Cell lysates were collected at various time points post-transfection, and activity was measured using the Cypridina Luciferase Glow Assay Kit (Thermo Scientific™, 16170) and the Gaussia Luciferase Glow Assay Kit (Thermo Scientific™, 16160). All assays were conducted in white 96-well plates using a plate reader (Biotek Synergy H1). Gaussia luciferase measurements were normalized by CLuc values to account for well-to-well transfection variability.

### Virus injection and stereotactic surgery

Eight-week-old wildtype mice were injected with AAVs (30 nl, titer > 10^12) at specific brain regions, including POA (AP 0.2 mm, ML 0.3 mm, DV −5.2 mm), DMH (AP −1.9 mm, ML 0.2 mm, DV −5.3 mm), PAG (AP −4.0 mm, ML 0.5 mm, DV −2.5 mm), Striatum (AP 1.2 mm, ML 2.5 mm, DV −3.5 mm), SC (AP −3.6 mm, ML 0.6 mm, DV −1.2 mm), and V1 (AP −3.5 mm, ML 2.5 mm, DV −1.2 mm), depending on the experimental objectives and the mouse strain. For optogenetic manipulation, an optic cannula (200 μm, RWD, Shenzhen, China) was stereotactically implanted following virus injection. The optic cannula was secured with dental cement (C & B Metabond Quick Adhesive Cement System, Parkell, Edgewood, NY).

After each experiment, brains were sectioned and imaged using an epifluorescence microscope (Olympus, Tokyo, Japan) to confirm viral expression and the location of the implanted cannula.

### Immunostaining, imaging, and quantification

Mice were anesthetized with 1–2% isoflurane and then perfused with phosphate-buffered saline (PBS, containing 137.0 mM NaCl, 2.7 mM KCl, 8.0 mM Na_2_HPO_4_, 1.8 mM KH_2_PO_4_, pH 7.4), followed by 4% paraformaldehyde (PFA). The brains were post-fixed in 4% PFA overnight at 4°C and then sectioned into 50-µm slices in PBS using a vibrating blade microtome (Leica VT1200S, Wetzlar, Germany).

For immunostaining, brain slices were blocked in blocking buffer (TBS with 0.3% Triton X-100, 10% normal donkey serum, and 2% bovine serum albumin) at room temperature for 2 hours, then incubated overnight at 4°C with primary antibodies against mCherry (rabbit anti-mCherry, 1:1000, Abcam, Cambridge, UK). This was followed by a 2-hour incubation with secondary antibodies (donkey anti-rabbit IgG conjugated with Alexa RRX, 1:1000, Jackson, West Grove, PA) at room temperature.

Tissue samples were mounted with Fluoromount G (Thermo-Fisher, Waltham, MA), and specimens were imaged using a confocal microscope (Leica SP8, Wetzlar, Germany) with a 40x objective after drying. Fluorescent-positive neurons were counted using *Cellpose* (48), a deep-learning-based algorithm for cell segmentation.

### RNAscope combined with immunofluorescence

Mouse brains were fixed in 4% paraformaldehyde (PFA) at 4 degrees overnight and cryoprotected in 30% sucrose at 4 degrees for 2 days. Brains were then embedded in OCT and frozen. Coronal sections (12 μm) were collected directly onto glass slides.

Multiplex fluorescent RNAscope was performed using the RNAscope Multiplex Fluorescent Reagent Kit v2 (ACDbio, 323100) according to the manufacturer’s protocol. Sections were hybridized with probes against tyrosine hydroxylase (ACDBio, 317621-C2) and tdTomato (ACDBio, 317041-C3). Following completion of the RNAscope amplification and fluorescent detection steps, but before DAPI staining and mounting, sections were processed for immunofluorescence staining using a rabbit anti-mCherry antibody (Abcam, ab167453; 1:1000 followed by donkey anti-rabbit IgG (H+L) Alexa Fluor 555 secondary antibody (Invitrogen, A31572; 1:1000). Sections were then mounted with Fluoromount-G with dapi.

### Fresh brain slice preparation

For patch-clamp recordings or Ca^2+^ imaging, coronal brain slices containing the DMH were prepared in chilled slicing solution (92.0 mM NMDG-Cl, 10.0 mM NaCl, 2.5 mM KCl, 0.5 mM CaCl_2_, 10.0 mM MgSO_4_, 20.0 mM HEPES, 1.25 mM NaH_2_PO_4_, 26.0 mM NaHCO_3_, 25.0 mM glucose, saturated with 95% O_2_ + 5% CO_2_). The 300-μm-thick brain slices were cut using a vibrating blade microtome (Leica VT1200S, Wetzlar, Germany). Non-targeted brain regions were trimmed away, and the slices were allowed to recover in the slicing solution at 35°C for 10 minutes. The slices were then transferred to a holding chamber containing artificial cerebrospinal fluid (ACSF: 116.0 mM NaCl, 2.5 mM KCl, 2.0 mM CaCl_2_, 1.0 mM MgSO_4_, 1.25 mM NaH_2_PO_4_, 26.0 mM NaHCO_3_, 25.0 mM glucose, saturated with 95% O_2_ + 5% CO_2_). The brain slices were incubated at room temperature for at least 1.5 hours before electrophysiological recordings or Ca^2+^imaging.

### Electrophysiology

Whole-cell patch-clamp recordings were performed in a submerged chamber superfused (2–3 ml/min) with oxygenated ACSF at room temperature. Neurons were visually identified using an upright microscope (Olympus BX51WI, Tokyo, Japan) equipped with DIC optics; filter sets for GFP or tdTomato visualization, and a CCD camera (FLIR, Wilsonville, OR). Slices were visualized with a 60× objective under infrared illumination. Both mScarlet-labeled and unlabeled neurons surrounded by ChR2-expressing axon terminals were selected for recordings.

Recordings were made using an Axon 700B amplifier, digitized with a Digidata 1550B, and recorded/analyzed using pClamp 11 software (Molecular Devices, San Jose, CA). Patch pipettes with resistances of 5–8 MΩ (Sutter, Novato, CA) were filled with internal solution containing 135.0 mM CsMeSO_4_, 1.0 mM EGTA, 10.0 mM HEPES, 3.8 MgCl_2_, 4.0 mM Na_2_-ATP, 0.4 mM Na_2_-GTP, and 10.0 mM Na_2_-phosphocreatine (adjusted to pH 7.3 with CsOH). The membrane potential was held at 0 mV to record outward inhibitory postsynaptic currents (IPSCs).

Photostimulation was performed using a 460 nm LED (Mshot, Guangzhou, China) controlled by pClamp 11 software. Light intensity was set to 100% (66.24mW) with a pulse duration of 5 ms at 1 Hz. Tetrodotoxin (TTX, 1 µM, Tocris, Bristol, UK) was bath-applied, followed by a combination of TTX and 4-aminopyridine (4-AP, 1 mM, MCE, Monmouth Junction, NJ). Access resistance was 15–30 MΩ and monitored throughout the experiment. Typically, only one neuron per slice was recorded. Data were filtered at 10 kHz and digitized at 20 kHz.

### Ca^2+^ imaging

Fresh brain slices were prepared as described above. After the recovery period, slices were mounted on a microscope stage perfused with ACSF. Confocal laser-scanning microscopy was performed using the Nikon A1R system equipped with a 40× 1.2 N.A. water-immersion objective (Nikon, Tokyo, Japan). The excitation wavelength was set to 470 nm, with emission detected between 500–550 nm. Imaging was performed at depths up to 60 μm from the tissue surface. Frame-scanning was conducted for 560 frames (600s total). A 635 nm laser (CNI, Changchun, China) was used for stimulation at 20 Hz for 120 seconds, followed by a bath application of high K^+^ (20 mM) in ACSF as a positive control. Regions of interest (ROIs) corresponding to individual neurons were manually selected. ΔF/F0 values were calculated as (F - F0) / F0, and the data were normalized to Z-scores using Matlab, which were subsequently used to create heatmaps.

### Chemogenetics

Following recovery from virus injection, a temperature probe (G2 E-Mitter, Starr Life Sciences, Oakmont, PA) was implanted into the mouse’s abdominal cavity. After an additional week of recovery, the mouse was single-housed in a temperature-controlled incubator (JNYQ, Ningbo, China) with a normal light schedule and an ambient temperature of 22°C for at least overnight to acclimate to the environment. Food and water were provided *ad libitum*. The cage was placed on an ER4000 Energizer/Receiver (Starr Life Sciences, Oakmont, PA), which powered the E-Mitter and recorded data.

On the experimental day, the mouse was injected intraperitoneally with deschloroclozapine (1 mg/kg body weight, MCE, Monmouth Junction, NJ) or vehicle (sterile saline). Body temperature and locomotor activity were continuously monitored using VitalView software (Starr Life Sciences, Oakmont, PA) at a sampling rate of one measurement per minute.

### Optogenetics

Animals that underwent virus injection and optic fiber implantation were subjected to both the open field test and the place preference test, with a 72-hour interval between the two behavioral sessions.

The open field test was conducted in a white box (60 cm × 60 cm × 40 cm, length × width × height), which was virtually divided into a central area (35 cm × 35 cm) and a peripheral area. For each test, the mouse was placed in the peripheral area, and a video camera recorded its locomotion for 20 minutes to measure the time spent in the central and peripheral areas. LED stimulation (635 nm, 10 Hz, 20 ms pulse duration) was applied continuously during the test. Mouse movement tracking and analysis were done using MouBeAT (49).

For the place preference test, a white behavioral box (40 cm × 20 cm × 20 cm) was divided into two chambers. During each trial, the mouse was initially placed in the non-stimulation chamber. LED stimulation (635nm, 10Hz, 20ms pulse duration) was delivered continuously once the animal entered the stimulation chamber and ceased when the animal exited. Each test session lasted 20 minutes, and behavioral data were recorded using a camera. The LED stimulation was automatically controlled in a closed-loop manner by the RWD Fiber Photometry System, which detected the animal’s location in real-time.

## Statistics

Statistical analyses were performed using Prism 10.1.1. Sample sizes were determined based on similar experiments in the field: 3–4 replicates for immunostaining, 10–20 replicates for electrophysiology, 4 replicates for Ca^2+^ imaging, 3 replicates for luciferase assay, and 6-7 replicates for animal behavior tests. All data are expressed as mean ± standard error of the mean (SEM). Statistical details, including the sample size (n) and p-values, are provided in the text or figure legends.

## Acknowledgments

We thank Drs. Dong Li, Wendy Yue, Kenichi Toma, and Xuaner Xiang for their constructive suggestions, and Ms. Xiaoqing Hao for assistance with stereotactic injections. We also thank the CIBR facilities for their support, including the Vector Core for virus packaging, the Laboratory Animal Resource Center for mouse husbandry, and the Optical Imaging Facility for technical assistance with imaging. We acknowledge support from the National Science and Technology Major Project (2025YFA1804600 to T.A.W.) and National Institutes of Health (U01NS136405 and R01EY030138 to X.D; R01NS116588 to L.Y.J.). L.Y.J. is HHMI Investigator Emerita.

## Author Contributions

Conceptualization, Formal Analysis, and Writing of Original Draft: C.C. and T.A.W.; Investigation: C.C., R.L., A.Y.L., X.C., J.H., S.G., S.C., and X.C.; Resources: W.Z., F.Z., and C.H.; Review & Editing of Manuscript, Supervision, and Funding Acquisition: X.D., L.Y.J., and T.A.W.

---

**Fig. S1.**
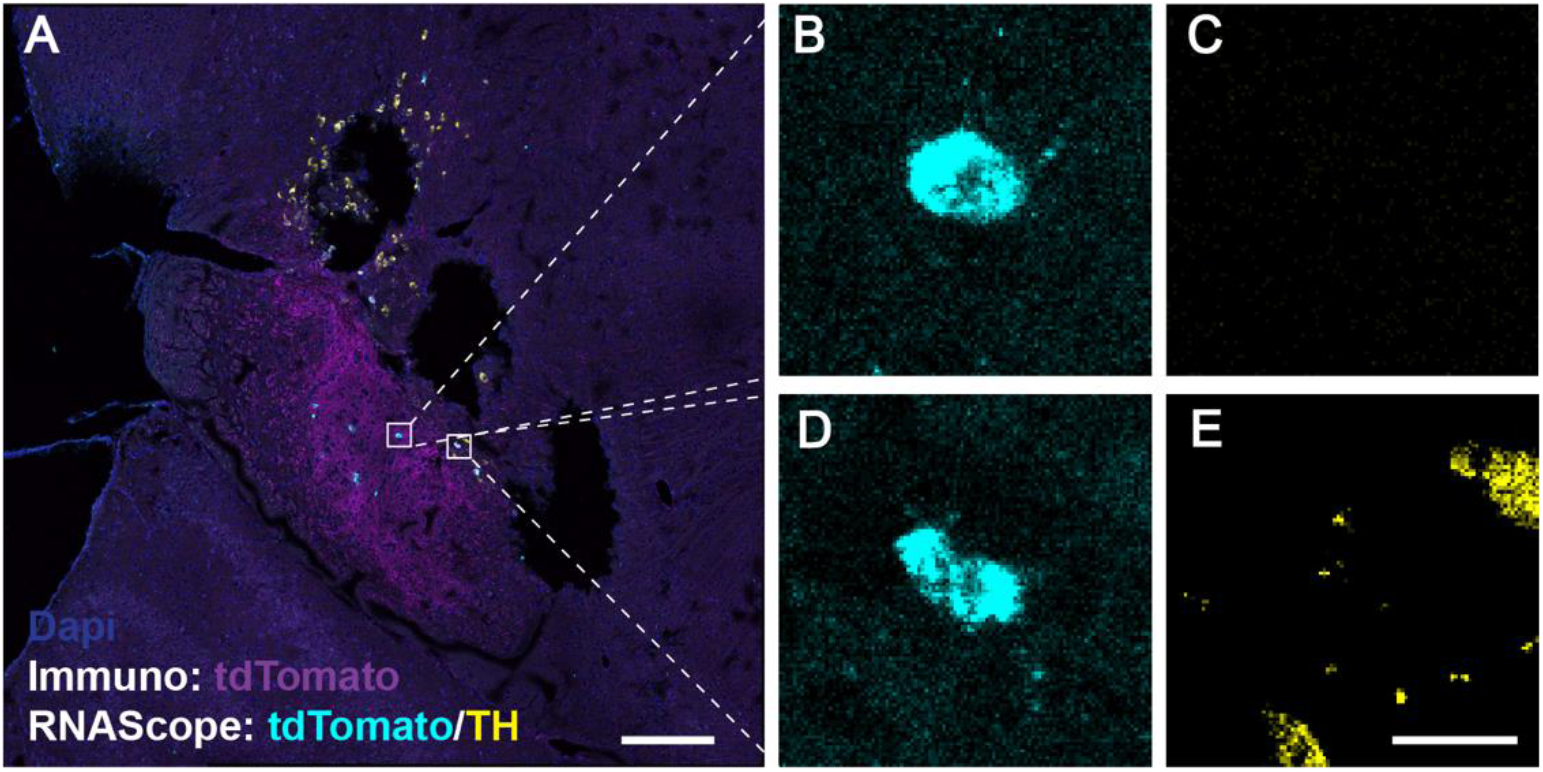
WTR-labeled neurons in the substantia nigra are TH negative **(A)** Representative image of the substantia nigra following WTR injection into the striatum. tdTomato immunostaining (magenta) shows WTR-associated labeling, while RNAscope was used to detect tdTomato mRNA (cyan) and TH mRNA (yellow). The boxed regions indicate representative tdTomato mRNA-positive cells shown at higher magnification in **B–E**.**(B-E)** High-magnification images of the boxed regions in a showing tdTomato mRNA-positive neuronal somata (cyan) and TH expression (yellow). The tdTomato mRNA-positive cells did not show detectable TH expression, indicating that the WTR labeled neurons are TH negative. Magenta, tdTomato immunostaining; cyan, tdTomato mRNA; yellow, TH mRNA; blue, DAPI. Scale bars, 250 μm (**A**) and 20 μm (**E**); the scale bar in **E** applies to **B–E**.

**Fig. S2.**
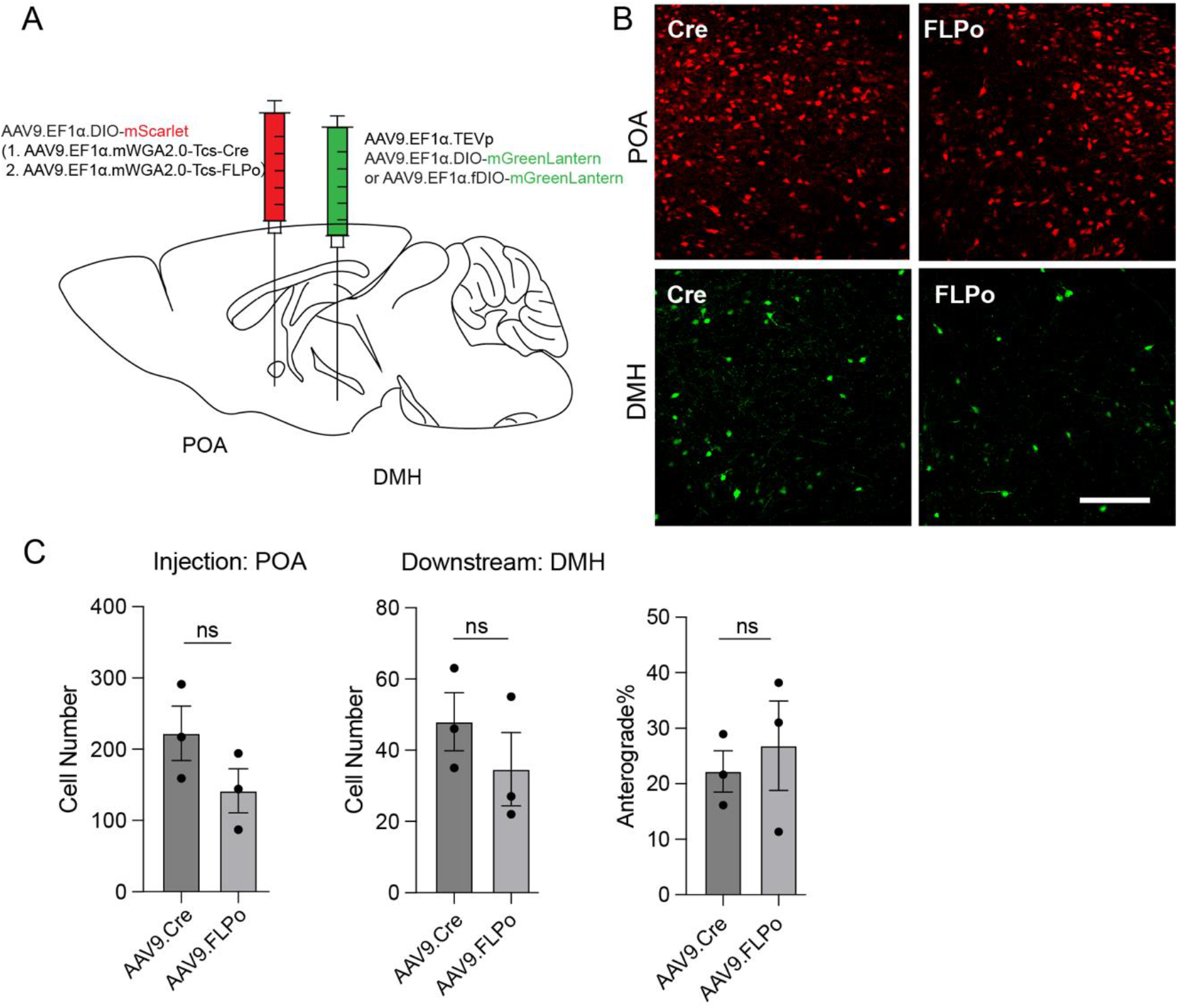
Cre and FLPo based WTR constructs show comparable anterograde tracing efficiency **(A)** Schematic illustration of the viral injection strategy used to compare mWGA2.0-TEVcs-Cre and mWGA2.0-TEVcs-Flpo in the POA–DMH pathway. WTR together with the corresponding recombinase dependent reporter was injected into POA, while TEVp and the corresponding Cre- or FLPo-dependent mGreenLantern reporter were injected into DMH. **(B)** Representative images showing reporter expression at the POA injection site and WTR-mediated labeling in downstream DMH neurons using Cre- or FLPo-based WTR constructs. **(C)** Quantification of labeled cells at the POA injection site, labeled downstream neurons in DMH, and normalized anterograde tracing efficiency. No significant difference was detected between Cre- and FLPo-based WTR constructs. n = 3 mice per group; ns, not significant. Data are shown as mean ± SEM. Scale bar, 100 μm.

**Fig. S3.**
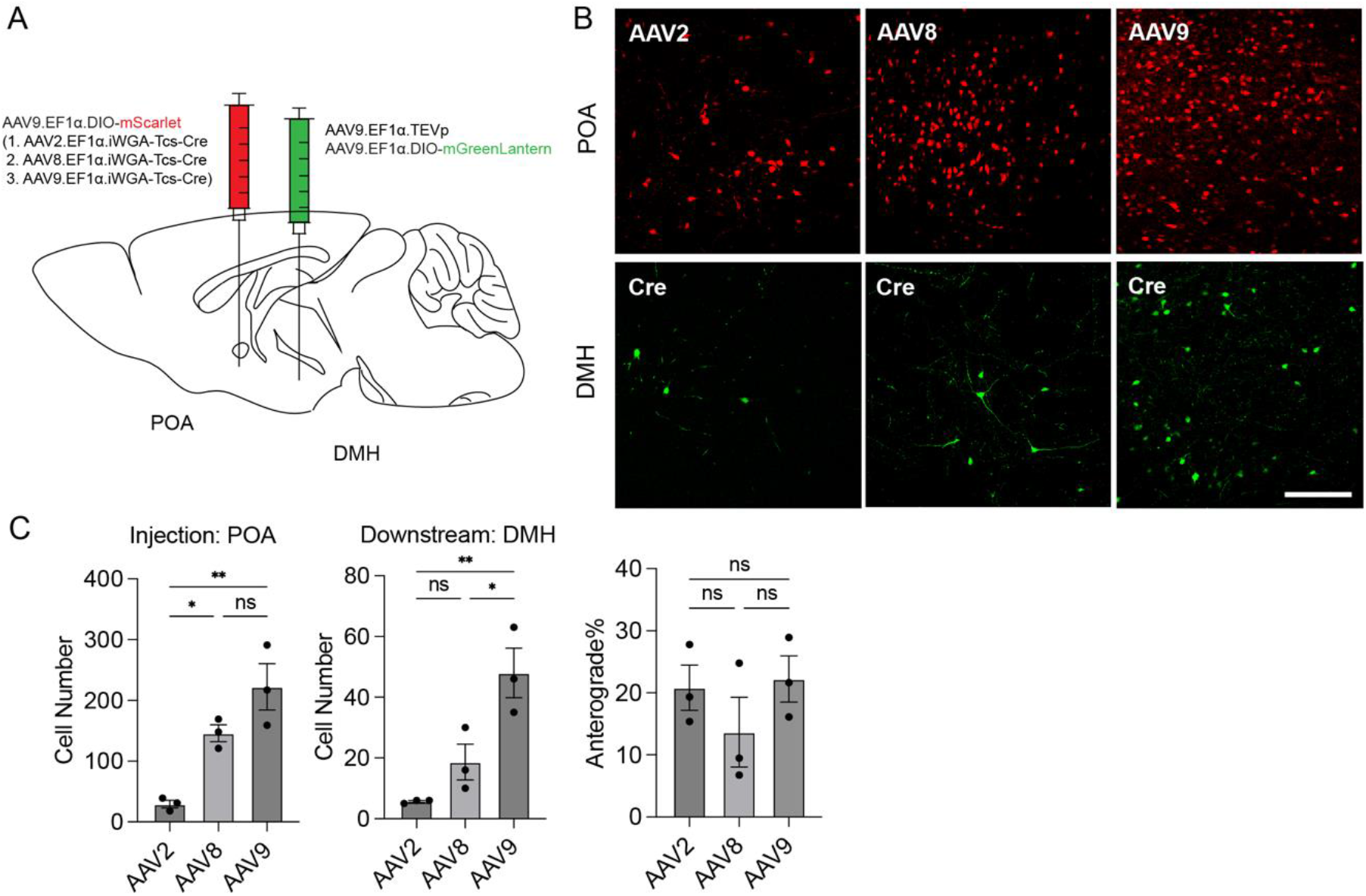
WTR anterograde efficiency is comparable across AAV serotypes **(A)** Viral tracer injection for anterograde tracing from POA to DMH using WTR packaged in different AAV serotypes. **(B)** Representative images showing fluorescent cells in POA and DMH. **(C)** Quantification of total labeled cells and anterograde tracing efficiency in POA and DMH across different groups. n = 3; *p < 0.05, **p < 0.01; one-way ANOVA followed by Tukey’s post-hoc test. Data are represented as mean ± SEM. Scale bar: 100 μm.

**Fig. S4:**
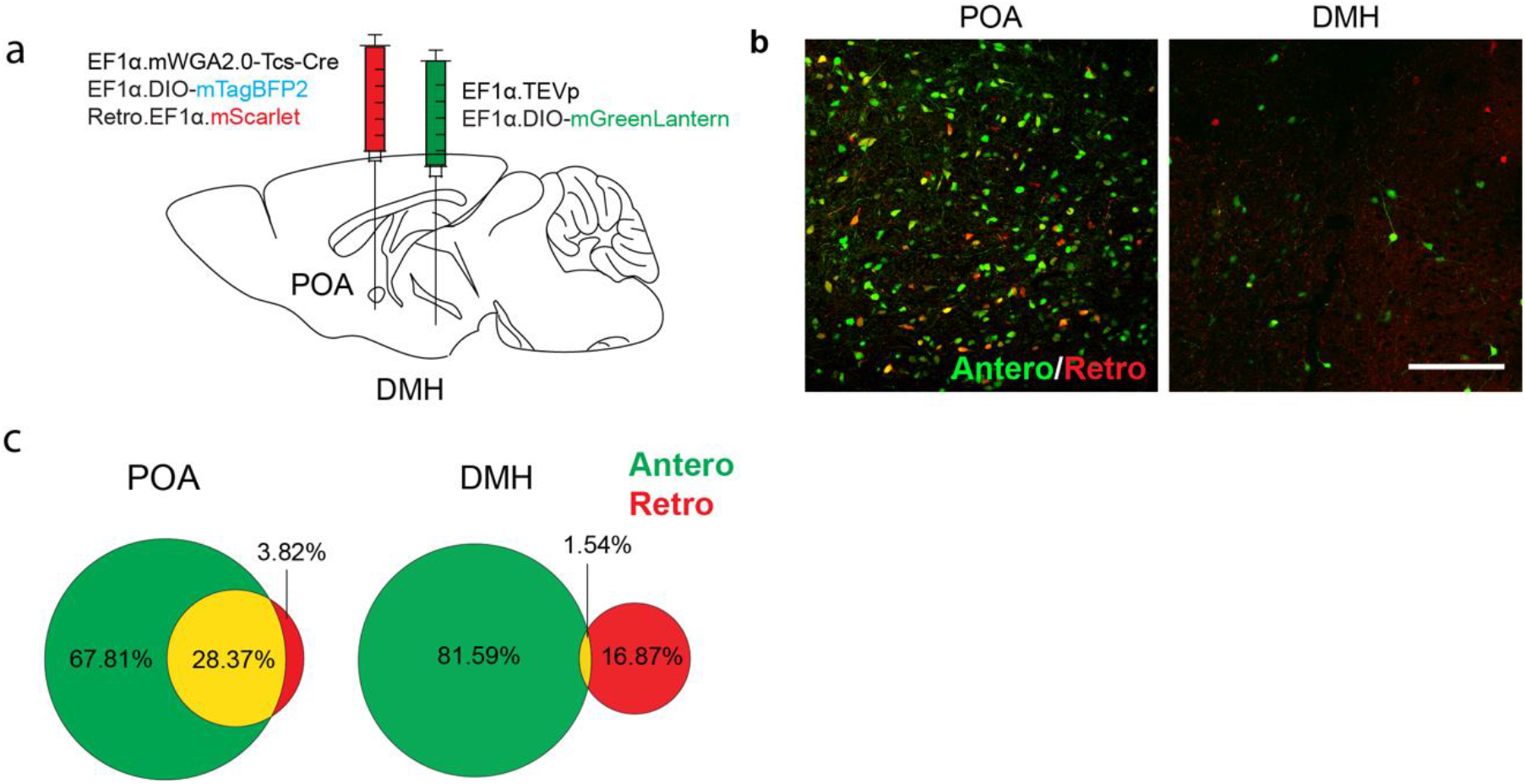
Combined anterograde and retrograde tracing distinguishes WTR labeled DMH neurons from POA projecting neurons **(A)** Viral injection setup for anterograde and retrograde tracings between POA and DMH. **(B)** Expression of mGreenLantern (indicating anterograde tracing) and mScarlet (indicating retrograde tracing) in DMH after injection of AAV2/9.iWGA-Tcs-Cre and AAVRetro.EF1α.mScarlet into POA (green, pseudo color of mTagBFP2 expression, indicating anterograde tracer transfection; red, mScarlet expression, indicating AAV.retro transfection). Scale bar: 100 μm. **(C)** Venn diagrams showing the percentage of neurons labeled exclusively by anterograde tracing (green), retrograde tracing (red), or both (yellow) in DMH. The transfection rate at the injection site is evaluated based on tracer expression in POA (anterograde: pseudo-green; retrograde: red). n=3 mice.

**Fig. S5.**
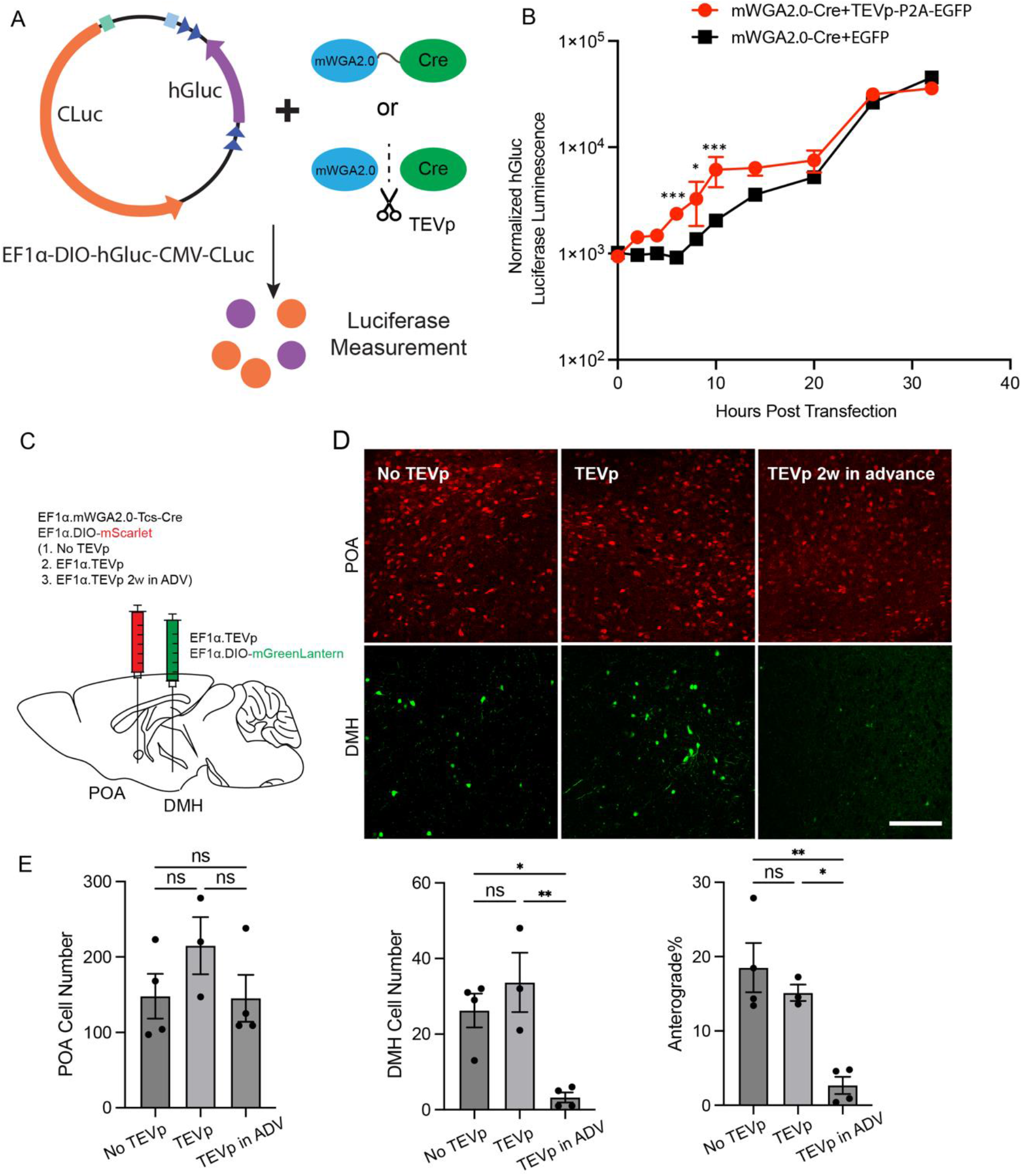
TEVp cleavage to release recombinase from the fusion protein enhances recombinase activity and reduces transfer of released recombinase **(A)** Schematic illustration of the dual-luciferase assay. HEK293T cells were transfected with the EF1α-DIO-hGluc-CMV-CLuc and mWGA2.0-Tcs-Cre plasmids, along with TEVp-P2A-EGFP or control EGFP plasmids. **(B)** Luciferase activity was measured at multiple time points post-transfection. n=3, Interaction (F (9, 40) =3.880, p=0.0013), Plasmid (F (1, 40) =40.95, p<0.0001), Time (F (9, 40)=166.9, p<0.0001). Plasmid effect at 6 hrs (p=0.0004), 8 hrs (p=0.019), and 10 hrs post-transfection (p=0.001) was significantly different, as assessed by two-way ANOVA on log-transformed data with Šídák’s multiple comparisons test. **(C)** Viral injection setup for anterograde tracing groups with or without TEVp in the POA: **No TEVp**: POA, AAV2/9.EF1α-mWGA2.0-TEVcs-Cre + AAV2/9.EF1α-DIO-mScarlet**; TEVp**: POA, AAV2/9.EF1α-mWGA2.0-TEVcs-Cre + AAV2/9.EF1α-DIO-mScarlet + AAV2/9.EF1α-DO-TEVp; **TEVp in Advance**: POA, AAV2/9.EF1α-DO-TEVp was injected 2 weeks prior to AAV2/9.EF1α-mWGA2.0-Tcs-Cre + AAV2/9.EF1α-DIO mScarlet; For all groups: DMH, AAV2/9.EF1α-DIO-mGreenLantern + AAV2/9.EF1α-TEVp. **(D)** Representative images showing fluorescent cells in the POA and DMH. Scale bar: 100 μm. **(E)** Quantification of total labeled cells and the percentage of anterograde labeling in the POA and DMH for the different groups. n=3–4; *p<0.05, **p<0.01, one-way ANOVA followed by Tukey’s post-hoc test. Data are presented as mean ± SEM.

**Fig. S6.**
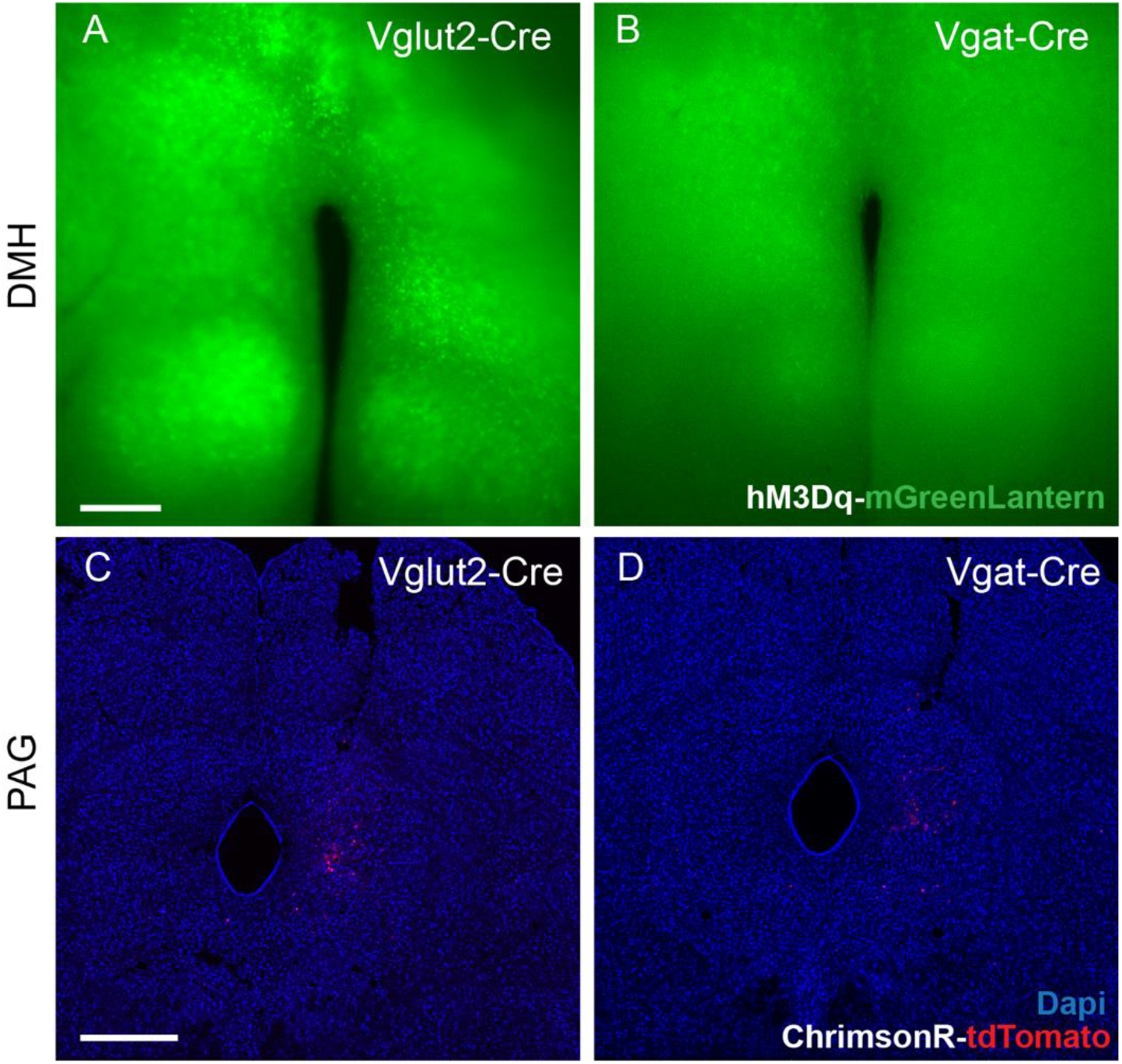
Expression of chemogenetic and optogenetic effectors in WTR labeled downstream neurons **(A,B)** Representative images showing hM3Dq-mGreenLantern expression in WTR-labeled DMH neurons downstream of POA Vglut2 neurons (**A**) or POA Vgat neurons (**B**) used for the body-temperature experiments in Fig. 5. **(C,D)** Representative images showing ChrimsonR-tdTomato expression in WTR-labeled PAG neurons downstream of POA Vglut2 neurons (**C**) or POA Vgat neurons (**D**) used for the behavioral experiments in **Fig. 5**. Green, hM3Dq-mGreenLantern; red, ChrimsonR-tdTomato; blue, DAPI. Scale bars, 250 μm (**A,C**); the scale bar in **A** applies to **A,B**, and the scale bar in **C** applies to **C,D**.

